# Limit of detection of Raman spectroscopy using polystyrene particles from 25 to 1000 nm in aqueous suspensions

**DOI:** 10.1101/2025.01.17.632088

**Authors:** Cindy Mayorga, Shreya Milind Athalye, Miad Boodaghidizaji, Neelesh Sarathy, Mahdi Hosseini, Arezoo Ardekani, Mohit S. Verma

## Abstract

Raman spectroscopy is an analytical method capable of detecting various microorganisms and small particles. Here, we used 25-1000 nm polystyrene particles in aqueous suspensions, which are comparable in size to viral particles and viral aggregates, to determine the limit of detection of a confocal Raman microscope. We collected Raman spectra using a 785 nm wavelength laser with a power of 300 mW and a 10 s exposure time, with a 5X objective lens. We detected the most prominent peak of the polystyrene particles at 1001 cm^-1^, corresponding to the ring breathing mode. We established the minimum and maximum limit of detection (LOD_min_ and LOD_max_) using a partial least squares (PLS) model. The LOD of the smallest size of 50 nm was identified as 7.47 × 10^12^ –7.64 × 10^12^ particle/mL, and for the largest size of 1000 nm, 6.09 × 10^8^ –6.24 × 10^8^ particle/mL. We demonstrated that Raman spectroscopy was non-destructive under these conditions by comparing the particle size before and after collecting Raman spectra using dynamic light scattering. Due to their size similarity to viral particles and viral aggregates, this systematic characterization of polystyrene particles provides detailed information on their Raman spectral signatures in aqueous suspensions. These findings establish a foundation for using Raman spectroscopy for the detection of small particles in aqueous suspensions and highlight its potential as a tool for real-time monitoring in vaccine manufacturing.

## INTRODUCTION

Vaccine manufacturing plays a critical role in controlling viral epidemics and pandemics.^1^ Vaccines serve as a primary defense mechanism against viral pathogens, and their rapid development and distribution are essential for effective epidemic and pandemic responses.^1,2^ Conventional methods for monitoring viral particles are limited for real-time monitoring of product quality and complicate efforts to meet the demands of vaccine production. This emphasizes the need for integrated technologies capable of real-time quality monitoring of viral particles and virus-like particles (VLPs) in vaccine manufacturing.

Conventional methods for monitoring viral particles, such as high-performance liquid chromatography (HPLC), dynamic light scattering (DLS), flow virometry, western blot, and transmission electron microscopy (TEM), are time-consuming, require sample preparation, and are usually used offline in the process.^3–6^ Spectroscopic techniques like near-infrared (NIR) spectroscopy, which is widely applied in pharmaceutical process monitoring,^7^ face challenges due to their high sensitivity to water absorption, limiting their use in aqueous systems.^8^ Due to these instrumental challenges, the applicability of these analytical tools for real-time monitoring in vaccine production is limited.

Vibrational spectroscopic techniques, such as Raman spectroscopy, have great potential for real-time monitoring. Some advantages of Raman spectroscopy encompass i) fast acquisition of results, ii) ease of use, iii) little to no sample preparation, and iv) suitability for organic and inorganic materials.^9,10^ Raman spectroscopy provides a chemical fingerprint through peak positions and relative peak intensities, enabling unique material identification and differentiation.^11^ Raman spectroscopy is a useful analytical tool for characterizing microorganisms for applications in agriculture, food production, pharmaceutical production, and environmental monitoring.^12–15^ It is one of many analytical techniques used to identify a variety of microorganisms^13,16,17^ and small particles.^18,19^ In biomanufacturing, Raman spectroscopy has been used to monitor mammalian cell cultures, pharmaceuticals, and monoclonal antibody manufacturing processes to ensure product quality.^5,20,21^ While Raman spectroscopy has demonstrated success in biomanufacturing processes, its applications in vaccine production are still emerging.

Prior studies have explored the utility of Raman spectroscopy for the identification and characterization of viral particles. For instance, Raman spectroscopy has been employed to detect falsified COVID-19 vaccines and predict the concentration of viral particles in ProQuad vaccine.^18,22^ Other works have explored its application for virus-like particles (VLPs). Raman spectroscopy has been utilized for a variety of applications in VLP research, including the quantification of VLP precipitation in Hepatitis B core antigen formulations, offline monitoring of SARS-CoV-2 VLPs in upstream production, detection and characterization of the spike protein in SARS-CoV-2 VLPs, and monitoring Zika VLPs throughout upstream production processes.^23–26^

Despite these applications of Raman spectroscopy, the impact of particle size and concentration on Raman spectra, especially for the sizes of viral particles and VLPs in aqueous suspension, remains poorly characterized. The size, size distribution, and concentration of small particles and molecules play an important role in optical properties, such as Raman spectroscopy.^27,28^ Yet, a systematic characterization of Raman spectra using well-defined particles in small sizes is lacking. Here, we used polystyrene particles in the range of 25-1000 nm as model particles to systematically study the effects of size and concentration on Raman signals in aqueous suspension. This size range of 25-1000 nm was selected to encompass dimensions relevant to viruses and VLPs, which typically range from 20-300 nm,^29^ with aggregates of viral particles extending to the upper limit of this range.^30–32^

In this study, we established a standardized method using Raman spectroscopy to detect model particles in the size range of viral particles, VLPs, and viral aggregates in aqueous suspensions. Using polystyrene particles as models, we systematically characterized particle sizes ranging from 25 to 1000 nm in aqueous suspensions using 96-well plates. We determined the minimum and maximum limits of detection for these model particles, identifying 50 nm as the smallest detectable particle size at concentrations between 7.47 × 10^12^ and 7.64 × 10^13^ particle/mL and 1000 nm as the largest detectable size at concentrations ranging between 6.09 × 10^8^ and 6.24 × 10^8^ particle/mL. While our work is based on model systems, these systems provide a comprehensive understanding of how small particles in aqueous suspension interact and influence Raman data acquisition. These results lay the groundwork for potential applications of Raman spectroscopy for real-time monitoring in vaccine production.

## EXPERIMENTAL SECTION

### Preparation of polystyrene particle concentrations

Polyspherex™ polystyrene particles (Phosphorex) suspended in 0.1% Tween 20 and 2mM NaNO_3_ (Millipore Sigma S2002-25G) in deionized water were used as model particles. The list of polystyrene sizes and concentrations is listed in Table 1. Polystyrene particles were vortexed before each use. Two-fold dilutions were prepared using 0.1% Tween 20 (Affymetrix USB CAS 9005-64-5) in water to obtain a total of 6 concentrations for all particle sizes.

**Table 1.**
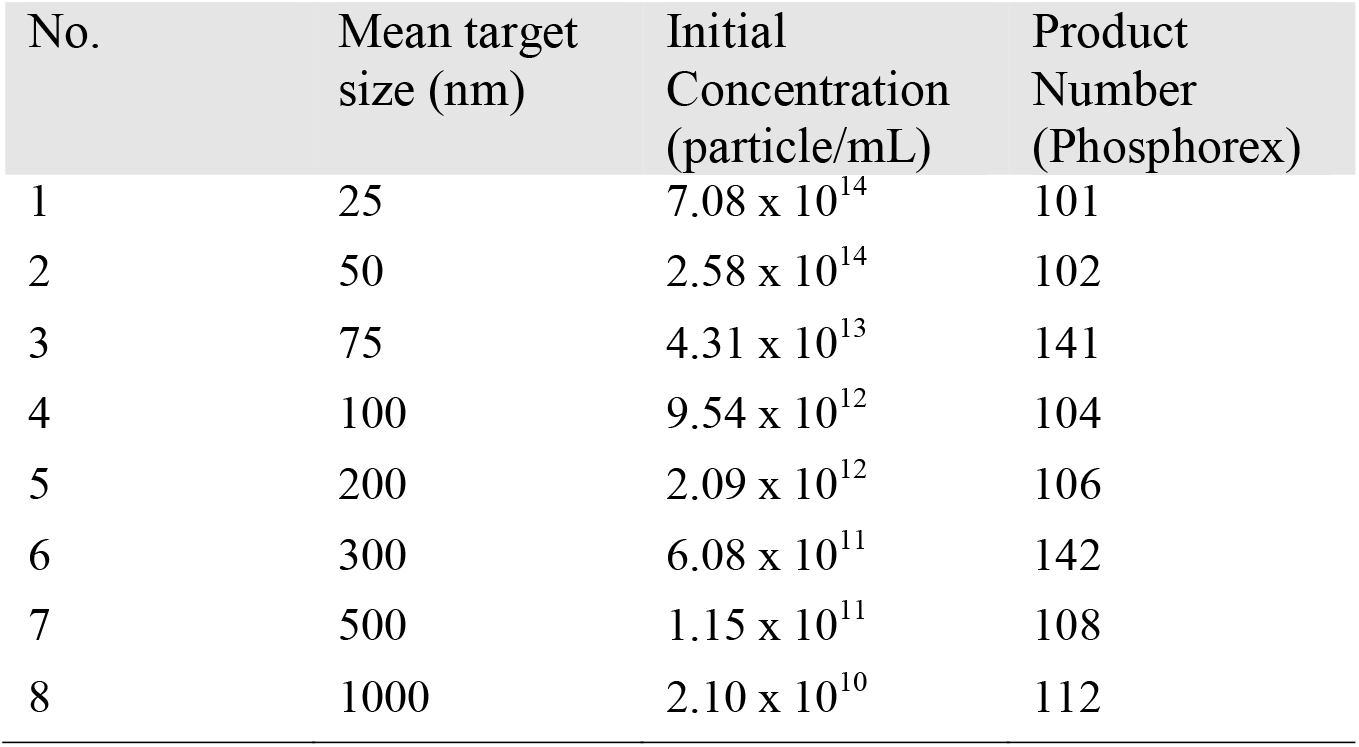
List of polystyrene particle sizes.

| No. | Mean target size (nm) | Initial Concentration (particle/mL) | Product Number (Phosphorex) |
| --- | --- | --- | --- |
| 1 | 25 | $7.08 \times 10^{14}$ | 101 |
| 2 | 50 | $2.58 \times 10^{14}$ | 102 |
| 3 | 75 | $4.31 \times 10^{13}$ | 141 |
| 4 | 100 | $9.54 \times 10^{12}$ | 104 |
| 5 | 200 | $2.09 \times 10^{12}$ | 106 |
| 6 | 300 | $6.08 \times 10^{11}$ | 142 |
| 7 | 500 | $1.15 \times 10^{11}$ | 108 |
| 8 | 1000 | $2.10 \times 10^{10}$ | 112 |

### Raman spectroscopy for the detection of polystyrene particles

Raman measurements were collected using a confocal Raman spectrometer (Renishaw inVia™ Qontor®) mounted on a microscope (Leica DM2700M). We used a 785-nm laser with 100% laser power (300 mW), 1200 lines/mm grating, and 10 s exposure time. The Raman data were collected in a spectrum ranged from 100-3200 cm^-1^ with a spectral resolution of 1 cm^-1^. The polystyrene particles were focused with a 5X objective lens. The 5X objective has the following optical properties: Numerical aperture = 0.12, Working distance = 14 mm, depth of focus = 109.02 µm, and laser spot = 8 µm. A 96-well plate (Corning™3635 Acrylic Clear Flat Bottom UV-Transparent Microplate) was used to take Raman spectra of polystyrene particle aqueous suspensions. All polystyrene particle sizes and concentrations were measured in individual wells filled with 350 μL of suspension volume. The 96-well plate was placed under the microscope objective to collect Raman spectra of an empty 96-well plate, 0.1% Tween 20 buffer, water and NaNO_3_ in 96-well plate, and polystyrene particle suspensions. Eight scans (technical replicates) were taken per sample on three different days for a total of 24 scans collected. Raw Raman spectral data was collected using WiRE™ 5.5 software.

### Raman data processing

Raman spectra data were baseline corrected using the adaptive iteratively reweighted Penalized Squares (airPLS) algorithm by Zhang et al.^33^ and Python script adapted by Lombardo^34^ of the University of Palermo. The Python script was modified to remove cosmic rays from Raman spectra and normalize the Raman data using standard normal variate (SNV) normalization. Python script was adapted to calculate the area under the curve (AUC) of characteristics peaks for polystyrene particles.^35^

### Partial Least Squares (PLS) regression and LODs range

Partial Least Squares (PLS) regression was employed to develop predictive models that define the correlation variability between Raman spectra intensity in the range of 600-1800 cm^-1^ and the size, concentration, technical replicates, and day of polystyrene particles. The data used for the PLS model were processed as described in the Raman data processing section. The prediction model was built and calibrated using a training set consisting of 90% of the measurement data, with the remaining 10% as a testing set for validation. The K-fold cross-validation method was used for model calibration and to prevent overfitting. The root-mean-square error of prediction (RMSEP) was used as an indicator of the quality of the predictive model. Variable importance in projection (VIP) scores were calculated to determine the importance of the predictors in the response variable. The minimum and maximum limits of detection (LOD_min_ and LOD_max_) were determined for each particle size following the method established by Allegrini and Olivieri.^36^ All the analyses for both PLS and LOD range were performed in Python (version 3.12.7). The Python script for these analyses is available online.^37^ PLS and LOD results are available in the supplementary material section.

### Dynamic Light Scattering (DLS) for particle size

Particle sizes were analyzed by dynamic light scattering (DLS) using a Zetasizer Nano ZS (Malvern Panalytical). Each size and concentration of polystyrene particles were analyzed before and after Raman data acquisition. The material setting was set to polystyrene latex particles, and the dispersant index was set to water in the zetasizer software (Version 7.13). The refractive index and viscosity of water were used as calculation parameters. A folded capillary zeta cell (DTS1070 Malvern Panalytical) was used to take the diameter measurements of polystyrene particles in aqueous suspensions. All experiments were performed at 25 °C using non-invasive backscatter detection at a scattering angle of 173° with an automatic measurement detection (set as 11 runs and 10 s run duration). The mean diameter was obtained by calculating the average of three measurements taken on three different days for nine readings per sample.

## RESULTS AND DISCUSSION

### Detection of polystyrene particles with Raman spectroscopy

We chose polystyrene particles in aqueous suspensions, in a range of 25-1000 nm in size, as models to determine the limits of detection in Raman spectroscopy. This size range was chosen because it aligns with the size of viral particles, virus-like particles (VLPs), and viral aggregates. These model particles allow us to directly investigate the effects of size and concentration on Raman spectra without the complexities associated with components, composition variability, and structure of viral particles, VLPs, and aggregates present in vaccines.

The Raman spectroscopy data for all polystyrene particles were collected using a confocal Raman spectroscopy with a 785-nm laser, 300 mW laser power, 10 s exposure time, 5X objective lens, and 1200 lines/mm grating. The 5X objective was chosen due to its larger depth of focus, which allows capturing Raman features of polystyrene particle suspensions in 96-well plate and enables collection of spectral information of the particle suspensions rather than the individual particles. We considered both size and concentration to determine the limits of detection in a confocal Raman spectroscopy. We collected eight technical replicates measured randomly at different points of the polystyrene particles in aqueous suspension and repeated the experiment across three different days. Figure 1 shows the average of 24 spectra, represented by a solid black line, with standard deviation depicted as a gray shaded area for water and the higher concentration of each polystyrene particle size detailed in Table 1. Figures S1 to S8 show the spectra of the 96-well plate, water, buffer, and all concentrations used for each polystyrene particle size. The Raman shift in the spectra were trimmed to the 400–3200 cm□^1^ region to highlight the most relevant molecular vibrations of the polystyrene particles.

**Figure 1.**
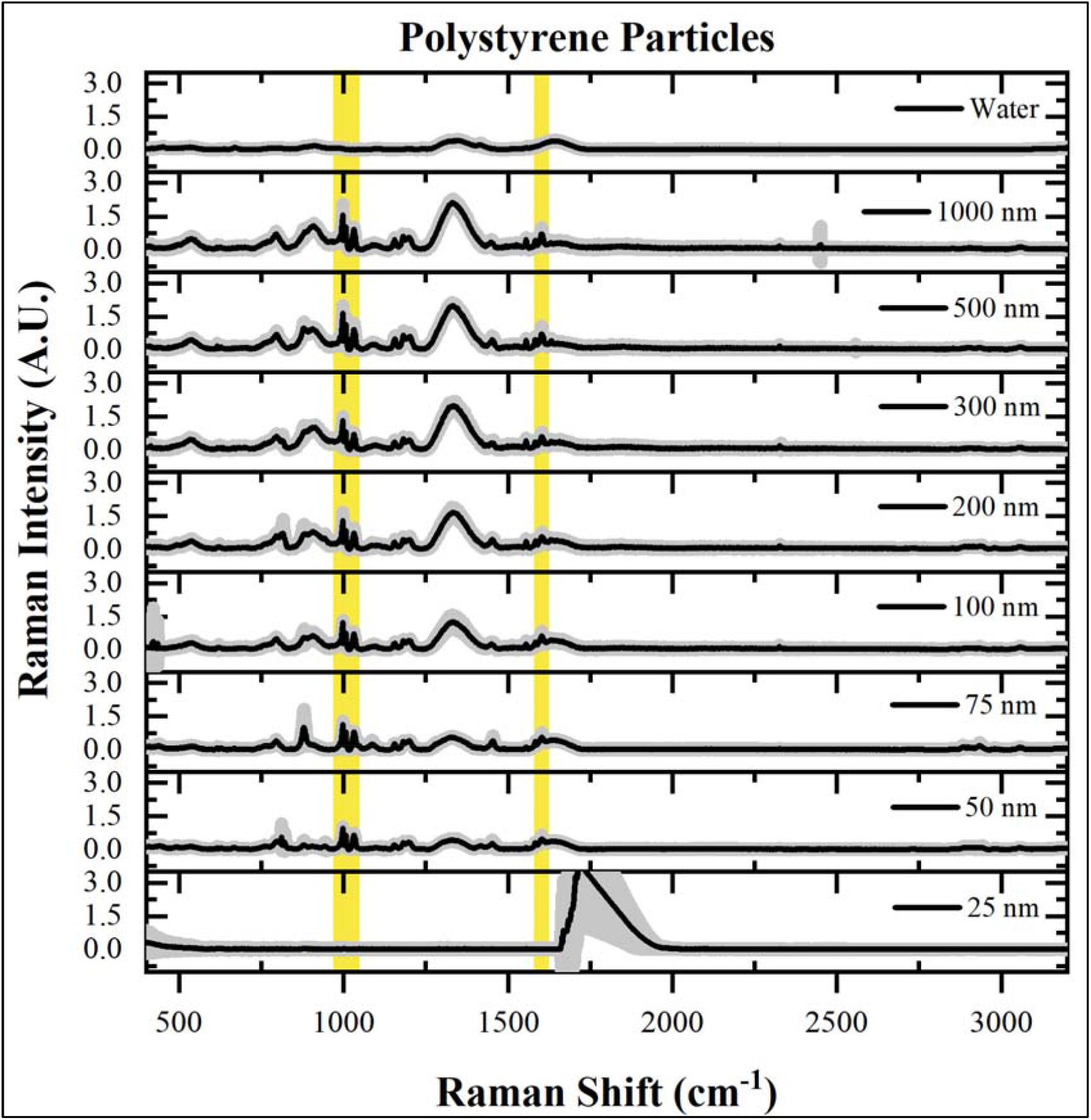
Raman spectra of water and polystyrene particles from 25 nm-1000 nm. Yellow shaded area represents the area of interest of the main peaks of polystyrene at 1001 cm^-1^ (990-1017 cm^-1^), 1031 cm^-1^ (1019-1047 cm^-1^) and 1602 cm^-1^ (1592-1615 cm^-1^). Raman spectra are averaged (n=24) and gray shaded area represents the standard deviation.

Raman spectra of 50 nm, 75 nm, 100 nm, 200 nm, 300 nm, 500 nm, and 1000 nm polystyrene particles showed polystyrene presence at 1001 cm^-1^,1031 cm^-1^ and 1602 cm^-1^ as prominent peaks, with the 1001 cm^-1^ peak being the most prominent. The 1001 cm^-1^ Raman peak corresponds to the ring breathing mode, the 1031 cm^-1^ Raman peak corresponds to C-H deformation in-plane and the 1602 cm^-1^ Raman peak to the ring-skeletal stretch.^38,39^ The Raman intensity of the 1001 cm^-1^,1031 cm^-1^ and 1602 cm^-1^ peaks decreased as the concentrations of polystyrene particles decreased. These peaks were absent in 96-well plate, water, 2 mM sodium azide, and 0.1% Tween 20 spectra, confirming they come from polystyrene particles. These 1001 cm^-1^, 1031 cm^-1^ and 1602 cm^-1^ peaks are present in all polystyrene particles except for the 25 nm size.

In the work by Mazilu et al.,^39^ Raman spectroscopy was used to detect a suspension of 2 μm size polystyrene beads with NIR dye. They detected peaks at 621, 1001, 1031, and 1602 cm^-1^ as the main peaks and confirmed our main detected peaks at 1001, 1031, and 1602 cm^-1^. Differences in the experimental setups and the smaller sizes of polystyrene beads used in this study may explain the absence of the 621 cm^-1^ peak observed in their work.

Polystyrene particles of 25 nm size showed charge-coupled device (CCD) saturation at higher concentrations and unsaturated Raman spectra at lower concentrations (Figure S1). Concentrations of polystyrene particles higher than 8.85 × 10^13^ particles/mL of 25 nm size can explain the CCD saturation. This is confirmed by the absence of CCD saturation at concentrations lower than 4.45 × 10^13^ particles/mL of 25 nm polystyrene particles. Although lower concentrations of 25 nm particles did not exhibit CCD saturation, they lacked any characteristic peaks of polystyrene particles.

We calculated the area under the curve (AUC) of the 1001 cm^-1^ prominent peak to quantify Raman signals of polystyrene particle concentrations. AUC considers how broad a peak is and provides a more accurate quantification when compared to peak intensity as a quantification method. The data pre-processing with baseline correction eliminates fluorescence, and normalization makes spectra measurements comparable across different days. Our data show (Figure 2) that AUC at 1001 cm^-1^ peak decreases as the concentration of particles decreases for all sizes, with a similar trend observed for smaller particles. For larger particles, the concentration of particles can be lower while still provide a Raman signal.

**Figure 2.**
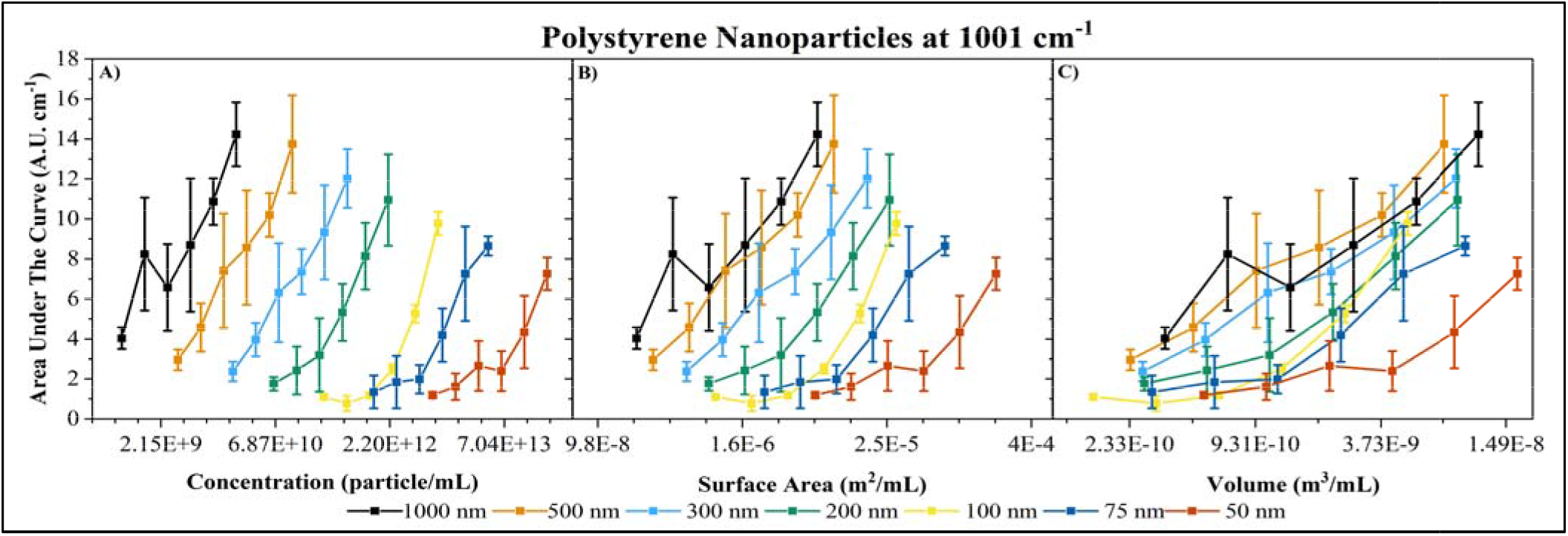
Area under the curve of polystyrene particles of sizes 50 nm, 75 nm, 100 nm, 200 nm, 300 nm, 500 nm, and 1000 nm at 1001 cm^-1^ peak in Raman shift as a function of A) concentration (particle/mL), B) surface area (m^2^/mL) and C) volume (m^3^/mL) (n=24).

The AUC is plotted as a function of concentration/volume, surface area/volume, and volume/volume (Figure 2). These three representations can provide insights into the properties and behavior of particles in liquid samples when interacting with the Raman laser. The higher the concentration of particles for a given size, the higher the Raman signal, and vice versa. As the size of particle decreases, their surface area per unit volume ratio increases. This higher surface area per unit volume exposes more functional groups that can exhibit Stokes and anti-Stokes scattering when exposed to the Raman laser. These data suggest that up to a certain limit, the higher the surface area per unit volume of particles, the higher the Raman signal. This aspect in the particles may enable the detection of particles as small as 50 nm. However, smaller particles exhibit more Rayleigh scattering and reduce the light that reaches the detector due to the limitation of the Raman spectroscopy instrument to detect smaller particles. Considering the surface area per unit volume, the total external surface area influences the interaction between particles of different sizes and the Raman laser. As the size of particles decreases, the Raman laser can interact with a larger surface area per unit volume, which helps to maintain a detectable Raman signal despite the reduction in signal strength due to the smaller size of particles.

### PLS regression for multivariate analysis and LOD estimation of Raman data

We have implemented a partial least square (PLS) as a multivariate analysis to estimate the limit of detection (LOD) of all polystyrene particle sizes. The PLS included the Raman spectra intensity in the range of 600-1800 cm^-1^ to include the relevant peaks of polystyrene. The overall PLS calibration model is used to estimate an LOD interval (LOD_min_ and LOD_max_).VIP scores from the PLS regression show that variables with scores above 1 are considered important in the model.^40^ In this analysis, size (VIP = 1.39) and concentration (VIP = 1.32) are the primary predictors influencing the Raman response of polystyrene particles, while day (VIP = 0.55) and technical replicates (VIP = 0.41) have a comparatively minor influence. Establishing an LOD_min_ and LOD_max_ for each particle size provides a detection range that shows the sensitivity and variation within the PLS model. Based on the methodology by Allegrini and Olivieri,^36^ this approach yields a robust LOD range that characterizes the detection capabilities of the PLS model across particle sizes. Values above LOD_max_ are considered reliable and confirm the presence of the analyte, while values below LOD_min_ are unreliable to distinguish the polystyrene particle from the background.

Figure 3 illustrates the LOD_min_ and LOD_max_ determined from the PLS model as a function of polystyrene particle size. Figure 3 was adjusted with an offset applied to the x-axis in the LOD_max_ to prevent overlapping. LOD_min_ and LOD_max_ values are presented in Table S3 for each particle size for reference. The LOD_min_ and LOD_max_ values decrease with increasing particle size, indicating that larger particles are detected at lower concentrations compared to smaller particles. This trend is expected, as small particles exhibit Rayleigh scattering, which occurs in particle sizes smaller than about 1/10 of the wavelength (i.e., about 80 nm). Rayleigh scattering occurs off the axis of incident light and dominates the loss for small particles. This phenomenon makes the detection of small particles in Raman spectroscopy more challenging. In contrast, Mie scattering occurs in larger particles, is more directional (along the incident light’s direction), and almost independent of particle size.^41–43^

**Figure 3.**
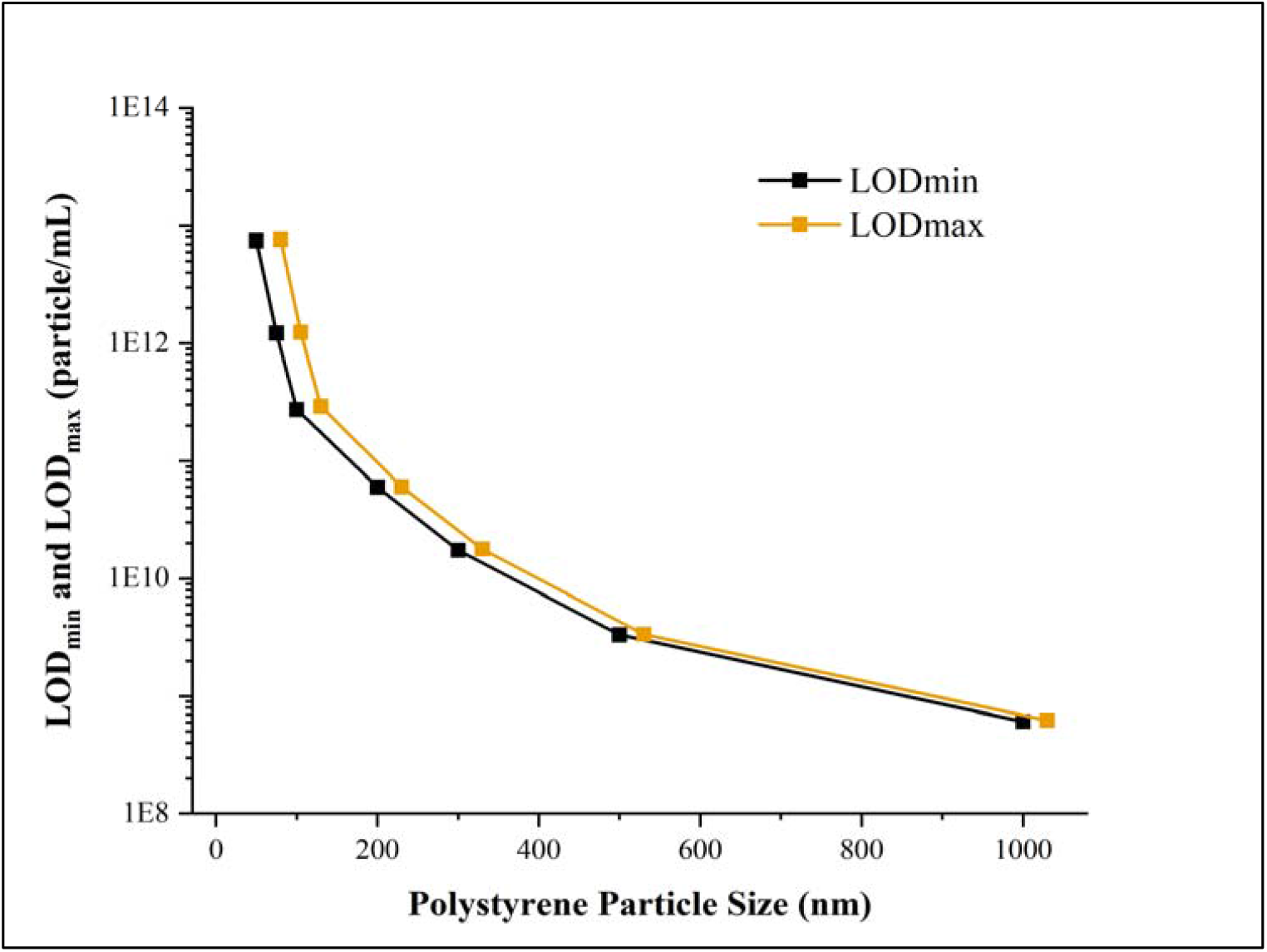
LOD_min_ and LOD_max_ as a function of polystyrene particle sizes. LOD_max_ data is offset along the x-axis to prevent overlapping.

For particles 100 nm and smaller, the LOD values are notably higher, which suggests reduced sensitivity of Raman spectroscopy in detecting smaller particles. This may be attributed to limitations in Raman signal strength and the PLS calibration model for these smaller particle sizes. This can explain the inability and challenges of Raman spectroscopy to detect 25 nm polystyrene particles. These findings suggest that Raman spectroscopy is more effective to detect larger particles and provides a guidance on the minimum particle size and concentration that can be reliably detected. The differences between the LOD_min_ and LOD_max_ values are relatively small, where LOD_max_ is slightly higher than LOD_min_. These small differences are probably due to consistent signal acquisition in the Raman spectroscopy setup established and the PLS calibration model. These narrow ranges reflect the reproducibility and precision of the Raman spectroscopy detection system and the reliable thresholds for each particle size.

### Effect of Raman laser power on particle size

We studied particle size before and after exposure to the Raman laser to determine potential particle damage. We presented the results as volume size distribution (Figures S9 to S16), which allowed us to evaluate changes in the size of particles in the samples. We observed some discrepancies in the size distribution of particles with concentrations of 200 nm, 300 nm, and 500 nm. These discrepancies were likely due to the turbid color at higher concentrations, which hindered light scattering. Our analysis of the volume size distribution data revealed that particle size remained unaffected before and after Raman laser acquisition, regardless of polystyrene particle size or concentration, when using a maximum laser power of 300 mW and 10 s of exposure time. It is likely that since the laser only focuses on a small volume within the sample during the acquisition of Raman spectra, the polystyrene particles in suspension remain undamaged by the laser.

## CONCLUSIONS

We employed polystyrene particles as model particles of viral particles, VLPs, and viral aggregates in the 25-1000 nm size range to evaluate the performance of a confocal Raman spectroscopy. The chosen experimental fixed setting with 785 nm wavelength, 5X objective lens, 10 s exposure time, and 300 mW laser power has the following four advantages: i) it can detect a wide range of particle sizes in liquid media using 96-well plates, ii) it allows to collect spectra of particles in their native state (or with a simple dilution), iii) it does not damage particles after several scans, and iv) it collects several spectra in less than 5 minutes.

The analysis of AUC enables the following two advantages: i) more accurate quantification of the signals in prominent peaks, and ii) visualization of the trend of particles at different sizes and concentrations. Moreover, the implementation of partial least squares regression allows the establishment of a robust LOD range for each particle size that demonstrates the sensitivity and variation of the model.

Two limitations of this study include: i) the characteristic Raman peak at 1001 cm^-1^ of polystyrene for 25 nm size could not be observed due to CCD saturation (this limitation for smaller particles could be overcome using a modified Raman setup that incorporates acoustic-wave-enhanced concentration or ultrasound particle manipulation^44–46^), and ii) the chosen experimental settings need further analysis and comparison with actual viral samples to validate our findings in a biological context. This is primarily because viral particles, VLPs, and viral aggregates do not always exhibit a spherical shape as polystyrene particles, which can affect Raman signal detection and data collection.

The Raman spectra data in this work were collected using a fixed setting and in a single confocal Raman microscope. We anticipate observing similar trends in the Raman spectra and quantification of AUC. We expect these similarities when employing similar sizes and concentrations across different Raman instruments and setups. This study aims to provide additional insights into the interaction of particle models in the size range of viral particles, VLPs, and viral aggregates in liquid samples, and how size and concentration can affect the performance in Raman spectra collection.

In future work, we aim to characterize and detect viral samples using Raman spectroscopy. We also aim to find the appropriate settings and experimental setup using Raman spectroscopy to detect particles as small as 25 nm in size.

## ASSOCIATED CONTENT

### Data availability statement

The data sets generated and/or analyzed during the current study are available in the Mendeley data repository at https://doi.org/10.17632/33wf5rtr4h.1.

### Supporting Information

Raman spectra for all particle sizes and concentrations, DLS graphs, the area under the curve for characteristic polystyrene particles, and PLS and LOD results.

## ACKNOWLEDGMENTS

This work was funded under a Project Award Agreement from the National Institute for Innovation in Manufacturing Biopharmaceuticals (NIIMBL) and financial assistance award 70NANB21H085 from the U.S. Department of Commerce, National Institute of Standards and Technology. We also thank Jiangshan Wang for reviewing this manuscript.

## Declaration of generative AI and AI-assisted technologies in the writing process

During the preparation of this work, the authors used ChatGPT for proofreading and clarification purposes. After using this tool/service, the authors reviewed and edited the content as needed and take full responsibility for the content of the publication.

## References

(1) Farlow, A.; Torreele, E.; Gray, G.; Ruxrungtham, K.; Rees, H.; Prasad, S.; Gomez, C.; Sall, A.; Magalhães, J.; Olliaro, P.; Terblanche, P. The Future of Epidemic and Pandemic Vaccines to Serve Global Public Health Needs. Vaccines 2023, 11 (3), 690. 10.3390/vaccines11030690.

(2) Rosa, S. S.; Prazeres, D. M. F.; Azevedo, A. M.; Marques, M. P. C. mRNA Vaccines Manufacturing: Challenges and Bottlenecks. Vaccine 2021, 39 (16), 2190–2200. 10.1016/j.vaccine.2021.03.038.

(3) Bhat, T.; Cao, A.; Yin, J. Virus-like Particles: Measures and Biological Functions. Viruses 2022, 14 (2), 383. 10.3390/v14020383.

(4) Neog, S.; Kumar, S.; Trivedi, V. Isolation and Characterization of Newcastle Disease Virus from Biological Fluids Using Column Chromatography. Biomed. Chromatogr. 2023, 37 (1), e5527. 10.1002/bmc.5527.

(5) Wasalathanthri, D. P.; Rehmann, M. S.; Song, Y.; Gu, Y.; Mi, L.; Shao, C.; Chemmalil, L.; Lee, J.; Ghose, S.; Borys, M. C.; Ding, J.; Li, Z. J. Technology Outlook for Real-Time Quality Attribute and Process Parameter Monitoring in Biopharmaceutical Development— A Review. Biotechnol. Bioeng. 2020, 117 (10), 3182–3198. 10.1002/bit.27461.

(6) Yang, N.; Zhang, J.; Xu, M.; Yi, J.; Wang, Z.; Wang, Y.; Chen, C. Virus-like Particles Vaccines Based on Glycoprotein E0 and E2 of Bovine Viral Diarrhea Virus Induce Humoral Responses. Front. Microbiol. 2022, 13, 1047001. 10.3389/fmicb.2022.1047001.

(7) Vovko, A. D.; Vrečer, F. Process Analytical Technology Tools for Process Control of Roller Compaction in Solid Pharmaceuticals Manufacturing. Acta Pharm. 2020, 70 (4), 443–463. 10.2478/acph-2020-0038.

(8) Wang, M.; An, H.; Cai, W.; Shao, X. Wavelet Transform Makes Water an Outstanding Near-Infrared Spectroscopic Probe. Chemosensors 2023, 11 (1), 37. 10.3390/chemosensors11010037.

(9) Schumacher, W.; Kühnert, M.; Rösch, P.; Popp, J. Identification and Classification of Organic and Inorganic Components of Particulate Matter via Raman Spectroscopy and Chemometric Approaches. J. Raman Spectrosc. 2011, 42 (3), 383–392. 10.1002/jrs.2702.

(10) Vankeirsbilck, T.; Vercauteren, A.; Baeyens, W.; Van der Weken, G.; Verpoort, F.; Vergote, G.; Remon, J. P. Applications of Raman Spectroscopy in Pharmaceutical Analysis. TrAC Trends Anal. Chem. 2002, 21 (12), 869–877. 10.1016/S0165-9936(02)01208-6.

(11) Schaldach, C. M.; Bench, G.; Deyoreo, J. J.; Esposito, T.; Fergenson, D. P.; Ferreira, J.; Gard, E.; Grant, P.; Hollars, C.; Horn, J.; Huser, T.; Kashgarian, M.; Knezovich, J.; Lane, S. M.; Malkin, A. J.; Pitesky, M.; Talley, C.; Tobias, H. J.; Woods, B.; Wu, K.-J.; Velsko, S. P. CHAPTER 13 – Non-DNA Methods for Biological Signatures. In Microbial Forensics; Breeze, R. G., Budowle, B., Schutzer, S. E., Eds.; Academic Press: Burlington, 2005; pp 251–294. 10.1016/B978-012088483-4/50016-0.

(12) Farber, C.; Sanchez, L.; Pant, S.; Scheuring, D.; Vales, I.; Mandadi, K.; Kurouski, D. Potential of Spatially Offset Raman Spectroscopy for Detection of Zebra Chip and Potato Virus Y Diseases of Potatoes (Solanum Tuberosum). ACS Agric. Sci. Technol. 2021, 1 (3), 211–221. 10.1021/acsagscitech.1c00024.

(13) Maruthamuthu, M. K.; Raffiee, A. H.; De Oliveira, D. M.; Ardekani, A. M.; Verma, M. S. Raman Spectra-Based Deep Learning: A Tool to Identify Microbial Contamination. MicrobiologyOpen 2020, 9 (11), e1122. 10.1002/mbo3.1122.

(14) Olaniyi, O. O.; Li, H.; Zhu, Y.; Cui, L. Metabolic Responses of Indigenous Bacteria in Chicken Faeces and Maggots to Multiple Antibiotics via Heavy Water Labeled Single-Cell Raman Spectroscopy. J. Environ. Sci. 2022, 113, 394–402. 10.1016/j.jes.2021.06.024.

(15) Vallejo-Pérez, M. R.; Sosa-Herrera, J. A.; Navarro-Contreras, H. R.; Álvarez-Preciado, L. G.; Rodríguez-Vázquez, Á. G.; Lara-Ávila, J. P. Raman Spectroscopy and Machine-Learning for Early Detection of Bacterial Canker of Tomato: The Asymptomatic Disease Condition. Plants 2021, 10 (8), 1542. 10.3390/plants10081542.

(16) de Siqueira e Oliveira, F. S.; da Silva, A. M.; Pacheco, M. T. T.; Giana, H. E.; Silveira, L. Biochemical Characterization of Pathogenic Bacterial Species Using Raman Spectroscopy and Discrimination Model Based on Selected Spectral Features. Lasers Med. Sci. 2021, 36 (2), 289–302. 10.1007/s10103-020-03028-9.

(17) Strycker, B. D.; Han, Z.; Duan, Z.; Commer, B.; Wang, K.; Shaw, B. D.; Sokolov, A. V.; Scully, M. O. Identification of Toxic Mold Species through Raman Spectroscopy of Fungal Conidia. PLOS ONE 2020, 15 (11), e0242361. 10.1371/journal.pone.0242361.

(18) Boodaghidizaji, M.; Milind Athalye, S.; Thakur, S.; Esmaili, E.; Verma, M. S.; Ardekani, A. M. Characterizing Viral Samples Using Machine Learning for Raman and Absorption Spectroscopy. MicrobiologyOpen 2022, 11 (6), e1336. 10.1002/mbo3.1336.

(19) Nava, V.; Frezzotti, M. L.; Leoni, B. Raman Spectroscopy for the Analysis of Microplastics in Aquatic Systems. Appl. Spectrosc. 2021, 75 (11), 1341–1357. 10.1177/00037028211043119.

(20) Domján, J.; Fricska, A.; Madarász, L.; Gyürkés, M.; Köte, Á.; Farkas, A.; Vass, P.; Fehér, C.; Horváth, B.; Könczöl, K.; Pataki, H.; Nagy, Z. K.; Marosi, G. J.; Hirsch, E. Raman-Based Dynamic Feeding Strategies Using Real-Time Glucose Concentration Monitoring System during Adalimumab Producing CHO Cell Cultivation. Biotechnol. Prog. 2020, 36 (6), e3052. 10.1002/btpr.3052.

(21) Maruthamuthu, M. K.; Rudge, S. R.; Ardekani, A. M.; Ladisch, M. R.; Verma, M. S. Process Analytical Technologies and Data Analytics for the Manufacture of Monoclonal Antibodies. Trends Biotechnol. 2020, 38 (10), 1169–1186. 10.1016/j.tibtech.2020.07.004.

(22) Mosca, S.; Lin, Q.; Stokes, R.; Bharucha, T.; Gangadharan, B.; Clarke, R.; Fernandez, L. G.; Deats, M.; Walsby-Tickle, J.; Arman, B. Y.; Chunekar, S. R.; Patil, K. D.; Gairola, S.; Van Assche, K.; Dunachie, S.; Merchant, H. A.; Kuwana, R.; Maes, A.; McCullagh, J.; Caillet, C.; Zitzmann, N.; Newton, P. N.; Matousek, P. Innovative Method for Rapid Detection of Falsified COVID-19 Vaccines through Unopened Vials Using Handheld Spatially Offset Raman Spectroscopy (SORS). Vaccine 2023, 41 (47), 6960–6968. 10.1016/j.vaccine.2023.10.012.

(23) Akdeniz, M.; Al-Shaebi, Z.; Altunbek, M.; Bayraktar, C.; Kayabolen, A.; Bagci-Onder, T.; Aydin, O. Characterization and Discrimination of Spike Protein in SARS-CoV-2 Virus-like Particles via Surface-Enhanced Raman Spectroscopy. Biotechnol. J. 2024, 19 (1), 2300191. 10.1002/biot.202300191.

(24) da Silva Cavalcante, P. E.; Públio Rabello, J.; Leme, J.; Aragão Tejo Dias, V.; Correia Barrence, F. A.; de Oliveira Guardalini, L. G.; Consoni Bernardino, T.; Almeida, S.; Tonso, A.; Attie Calil Jorge, S.; Fernández Núñez, E. G. Raman Laser Intensity and Sample Clarification on Biochemical Monitoring over Zika-VLP Upstream Stages. Biochem. Biophys. Res. Commun. 2024, 733, 150671. 10.1016/j.bbrc.2024.150671.

(25) Dietrich, A.; Schiemer, R.; Kurmann, J.; Zhang, S.; Hubbuch, J. Raman-Based PAT for VLP Precipitation: Systematic Data Diversification and Preprocessing Pipeline Identification. Front. Bioeng. Biotechnol. 2024, 12, 1399938. 10.3389/fbioe.2024.1399938.

(26) Moura Dias, F.; Públio Rabello, J.; Oliveira Guardalini, L. G.; Leme, J.; Consoni Bernardino, T.; Pires, L.; Mendonça, M.; Tonso, A.; Attie Calil Jorge, S.; Fernández Núñez, E. G. Laser Wavelength and Sample Conditioning Effects on Biochemical Monitoring of SARS-CoV-2 VLP Production Upstream Stage by Raman Spectroscopy. Biochem. Eng. J. 2024, 211, 109441. 10.1016/j.bej.2024.109441.

(27) Chander, N.; Khan, A. F.; Thouti, E.; Sardana, S. K.; Chandrasekhar, P. S.; Dutta, V.; Komarala, V. K. Size and Concentration Effects of Gold Nanoparticles on Optical and Electrical Properties of Plasmonic Dye Sensitized Solar Cells. Sol. Energy 2014, 109, 11–23. 10.1016/j.solener.2014.08.011.

(28) Sarra, A.; Stanchieri, G. D. P.; De Marcellis, A.; Bordi, F.; Postorino, P.; Palange, E. Laser Transmission Spectroscopy Based on Tunable-Gain Dual-Channel Dual-Phase LIA for Biological Nanoparticles Characterization. IEEE Trans. Biomed. Circuits Syst. 2021, 15 (1), 177–187. 10.1109/TBCAS.2021.3060569.

(29) Yu, H.; Afshar-Mohajer, N.; Theodore, A. D.; Lednicky, J. A.; Fan, Z. H.; Wu, C.-Y. An Efficient Virus Aerosol Sampler Enabled by Adiabatic Expansion. J. Aerosol Sci. 2018, 117, 74–84. 10.1016/j.jaerosci.2018.01.001.

(30) Gerba, C. P.; Betancourt, W. Q. Viral Aggregation: Impact on Virus Behavior in the Environment. Environ. Sci. Technol. 2017, 51 (13), 7318–7325. 10.1021/acs.est.6b05835.

(31) Petersen, J. D.; Lu, J.; Fitzgerald, W.; Zhou, F.; Blank, P. S.; Matthies, D.; Zimmerberg, J. Unique Aggregation of Retroviral Particles Pseudotyped with the Delta Variant SARS-CoV-2 Spike Protein. Viruses 2022, 14 (5), 1024. 10.3390/v14051024.

(32) Gupta, D.; Parthasarathy, H.; Sah, V.; Tandel, D.; Vedagiri, D.; Reddy, S.; Harshan, K. H. Inactivation of SARS-CoV-2 by β-Propiolactone Causes Aggregation of Viral Particles and Loss of Antigenic Potential. Virus Res. 2021, 305, 198555. 10.1016/j.virusres.2021.198555.

(33) Zhang, Z.-M.; Chen, S.; Liang, Y.-Z. Baseline Correction Using Adaptive Iteratively Reweighted Penalized Least Squares. Analyst 2010, 135 (5), 1138–1146. 10.1039/B922045C.

(34) Lombardo, R. airPLS.py. GitHub. https://github.com/zmzhang/airPLS/blob/master/README.md (accessed 2023-07-25).

(35) Ardekani, A. ArezooArdekani/Raman_spcetroscopy_baseline_correction, 2023. https://github.com/ArezooArdekani/Raman_spcetroscopy_baseline_correction (accessed 2023-11-07).

(36) Allegrini, F.; Olivieri, A. C. IUPAC-Consistent Approach to the Limit of Detection in Partial Least-Squares Calibration. Anal. Chem. 2014, 86 (15), 7858–7866. 10.1021/ac501786u.

(37) Verma Lab. VermaLab/Raman-PLSR: Performing Partial Least Squares Regression and Calculating Limit of Detection for Raman Spectroscopy, 2024. https://github.itap.purdue.edu/VermaLab/Raman-PLSR (accessed 2024-12-10).

(38) Bridges, T. E.; Houlne, M. P.; Harris, J. M. Spatially Resolved Analysis of Small Particles by Confocal Raman Microscopy: Depth Profiling and Optical Trapping. Anal. Chem. 2004, 76 (3), 576–584. 10.1021/ac034969s.

(39) Mazilu, M.; Luca, A. C. D.; Riches, A.; Herrington, C. S.; Dholakia, K. Optimal Algorithm for Fluorescence Suppression of Modulated Raman Spectroscopy. Opt. Express 2010, 18 (11), 11382–11395. 10.1364/OE.18.011382.

(40) Akarachantachote, N.; Chadcham, S.; Saithanu, K. CUTOFF THRESHOLD OF VARIABLE IMPORTANCE IN PROJECTION FOR VARIABLE SELECTION. Int. J. Pure Apllied Math. 2014, 94 (3). 10.12732/ijpam.v94i3.2.

(41) Gu, M.; Gan, X.; Deng, X. Scattering of Light by Small Particles. In Microscopic Imaging Through Turbid Media: Monte Carlo Modeling and Applications; Gu, M., Gan, X., Deng, X., Eds.; Springer Berlin Heidelberg: Berlin, Heidelberg, 2015; pp 15–23. 10.1007/978-3-662-46397-0_2.

(42) Hong, S.-H.; Winter, J. Size Dependence of Optical Properties and Internal Structure of Plasma Grown Carbonaceous Nanoparticles Studied by in Situ Rayleigh-Mie Scattering Ellipsometry. J. Appl. Phys. 2006, 100 (6), 064303. 10.1063/1.2338132.

(43) Lockwood, D. J. Rayleigh and Mie Scattering. In Encyclopedia of Color Science and Technology; Luo, M. R., Ed.; Springer New York: New York, NY, 2016; pp 1097–1107. 10.1007/978-1-4419-8071-7_218.

(44) Wheaton, S.; Gelfand, R. M.; Gordon, R. Probing the Raman-Active Acoustic Vibrations of Nanoparticles with Extraordinary Spectral Resolution. Nat. Photonics 2015, 9 (1), 68–72. 10.1038/nphoton.2014.283.

(45) Wieland, K.; Tauber, S.; Gasser, C.; Rettenbacher, L. A.; Lux, L.; Radel, S.; Lendl, B. In-Line Ultrasound-Enhanced Raman Spectroscopy Allows for Highly Sensitive Analysis with Improved Selectivity in Suspensions. Anal. Chem. 2019, 91 (22), 14231–14238. 10.1021/acs.analchem.9b01105.

(46) Kim, T.; Esmaili, E.; Athalye, S. M.; Matos, T.; Hosseini, M.; Verma, M. S.; Ardekani, A. M. Acoustofluidic Device Focusing Viral Nanoparticles for Raman Microscopy. Sens. Actuators B Chem. 2024, 420, 136438. 10.1016/j.snb.2024.136438.

