## Supporting Information for "Limit of detection of Raman spectroscopy using polystyrene particles from 25 to 1000 nm in aqueous suspensions"

**for**

Table of Contents for Supporting Information

### Summary of Raman spectroscopy and polystyrene particle studies

**Table S1**: Applications of different Raman spectroscopy methods on a variety of polystyrene particles.

| Nanoparticle material | Nanoparticle size | Raman Type | Wavelength | Laser power | Study Purpose | Conclusion | Reference |
| --- | --- | --- | --- | --- | --- | --- | --- |
| Polystyrene nanosphere and titanium dioxide nanosphere | 20 nm | Extraordinary acoustic Raman (EAR) spectroscopy | 853 nm | < 20 mW^[1]^ | Determine Raman-active acoustic modes from single nanoparticles in the 0.1-10 cm^-1^ range | EAR demonstrated a spectral resolution close to 3 orders of magnitude compared to typical Raman spectroscopy | ^1^ |
| Polystyrene nanoparticles in water | 60 nm and 120 nm | Fiber-enhanced system with Raman spectroscopy (FERS) | 532 nm | 1.7 mW and 50 mW | Chemical identification and concentration of polystyrene nanoparticles in aqueous samples | Raman signal was undetected for 121 nm size and within 0.92x10^12^ -3.72x10^12^ particles/mL for 63 nm size. | ^2^ |
| Polystyrene nanoparticles | 23-60 nm | Raman spectroscopy | 785 nm | 400 mW | Particle size prediction | Prediction of particle size with hybrid model R^2^=0.99 and with  data-driven partial least squares (PLS) regression R^2^=0.78-0.83. | ^3^ |
| Fluorescent polystyrene nanoparticles | 50 nm and 100 nm | Confocal Raman spectroscopy | 785 nm | 300 mW | Nanoparticle localization in cells and subcellular environment | Raman spectroscopy combined with K-means clustering effectively detects and locates particles in cells. | ^4^ |
| Polystyrene nanoparticles | 20 nm | Confocal Raman spectroscopy with optical trapping | 785 nm | 20 mW | Detection of polystyrene nanoparticles in plasmonic nanopores | Raman signals of low concentration of polystyrene nanoparticles are only possible with up-concentration in the nanopore. | ^5^ |
| Polystyrene nanoparticles | 20 nm | Surface-enhanced Raman spectroscopy (SERS) with optical trapping | 785 nm | 6, 10, 13 and 16 mW | Detection of polystyrene nanoparticles in plasmonic nanopores | A combination of SERS and optical trapping can detect small particles up to ~10 s with no particle damage. | ^6^ |
| Polystyrene nanoparticles | 200 nm - 10 µm | Inverted confocal Raman spectroscopy | 647.1 nm | 25 mW | Spatially resolved in the detection of different polystyrene nanoparticle sizes optically trapped | The confocal volume in this study was found to be ~1.3 fL. | ^7^ |
| Polystyrene particles | 25 nm - 1000 nm | Confocal Raman spectroscopy | 785 nm | 300 mW | Characterization of polystyrene particles as model systems in the size range of viral particles and aggregates | Raman instrument with fixed settings can detect nanoparticles as small as 50 nm at an LOD between 7.47 x 10^12^ and 7.64 x 10^13^ particle/mL. | This study |

### Supporting Figures

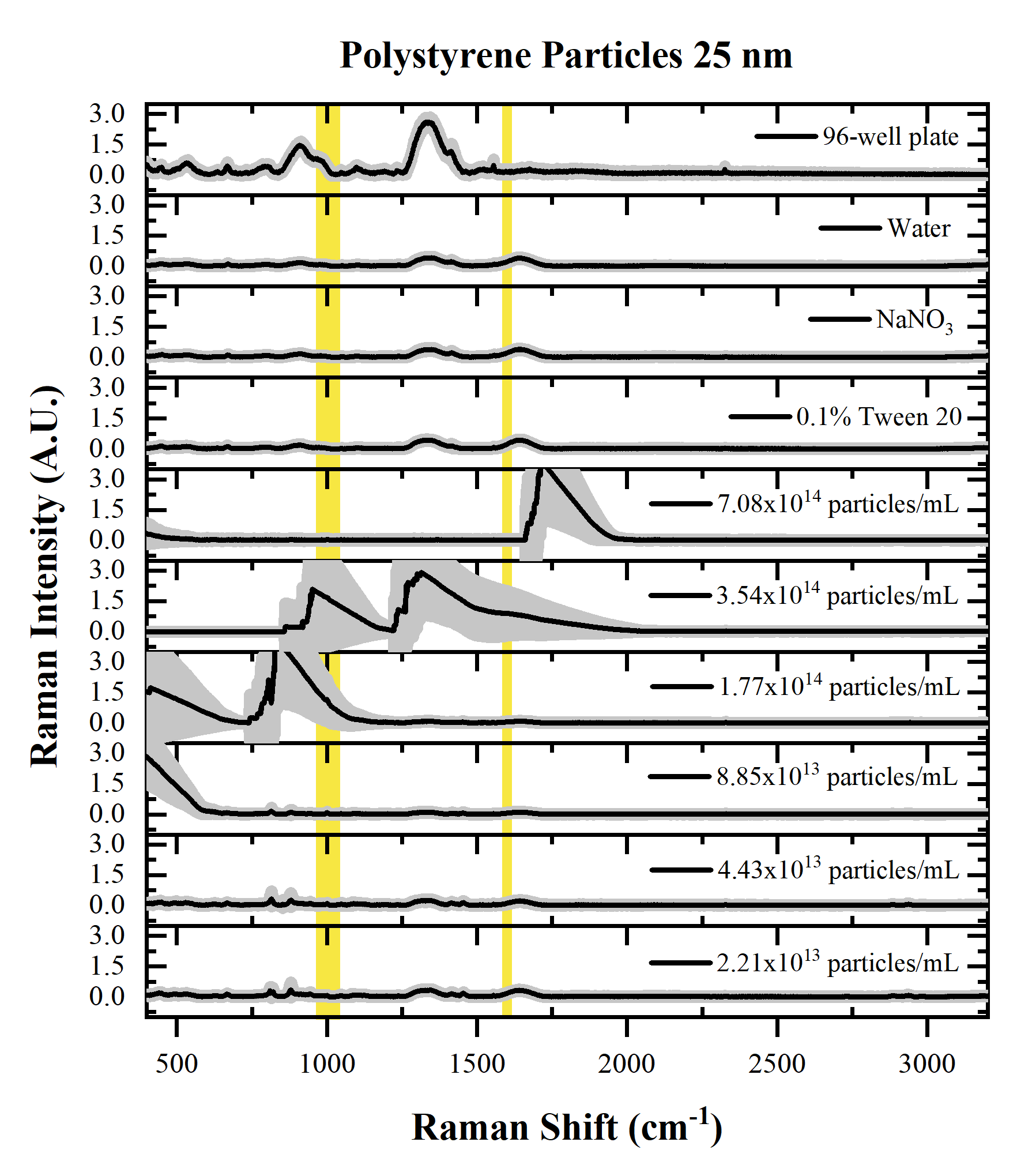

**Figure S1**: Raman spectra of 25 nm polystyrene particles. Yellow shaded area represents the area of interest of the main peaks of polystyrene at 1001 cm^-1^ (990-1017 cm^-1^), 1031 cm^-1^ (1019-1047 cm^-1^) and 1602 cm^-1^ (1592-1615 cm^-1^). Raman spectra are averaged (n=24) and gray shaded area represents the standard deviation.

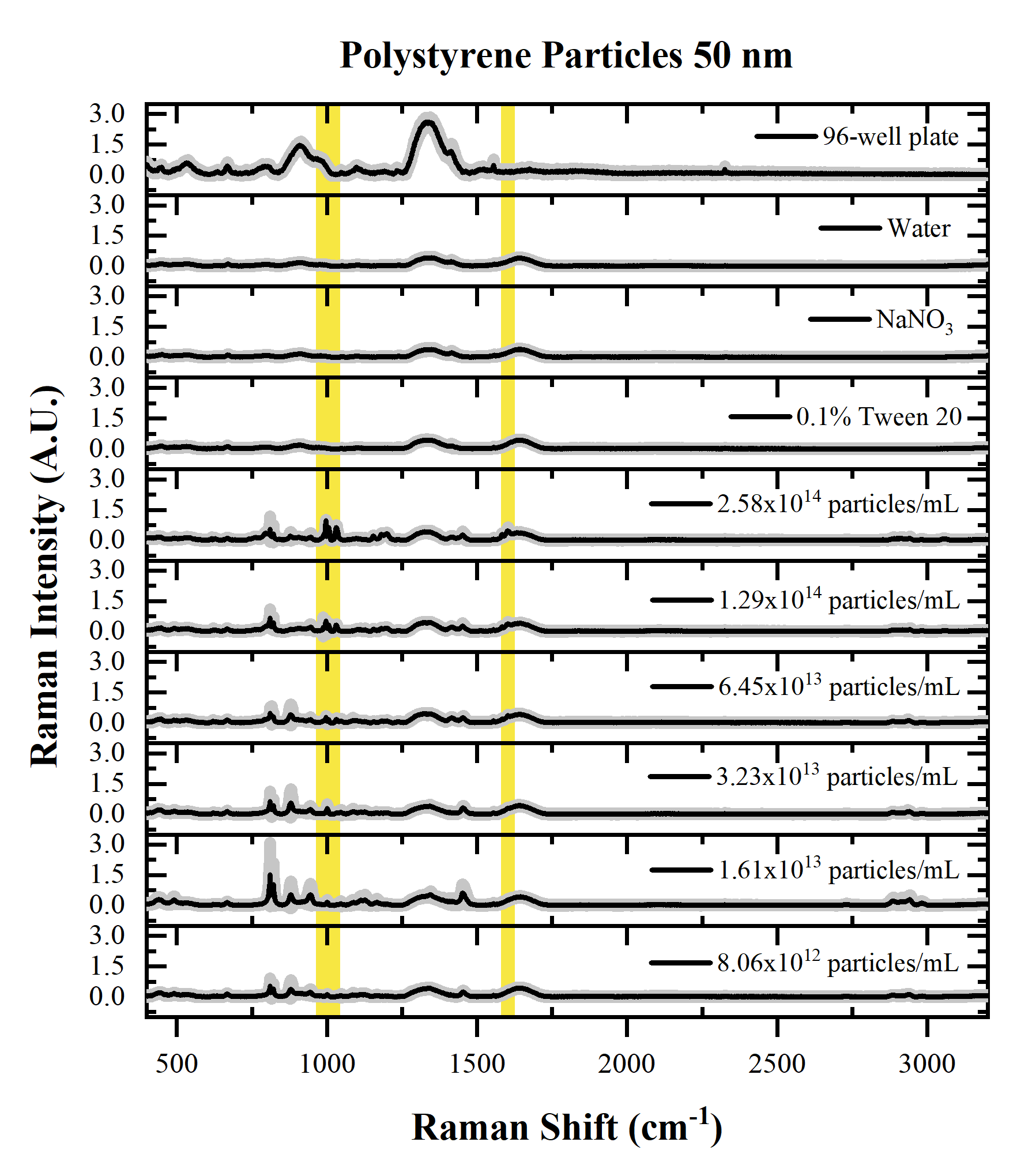

**Figure S2**: Raman spectra of 50 nm polystyrene particles. Yellow shaded area represents the area of interest of the main peaks of polystyrene at 1001 cm^-1^ (990-1017 cm^-1^), 1031 cm^-1^ (1019-1047 cm^-1^) and 1602 cm^-1^ (1592-1615 cm^-1^). Raman spectra are averaged (n=24) and gray shaded area represents the standard deviation.

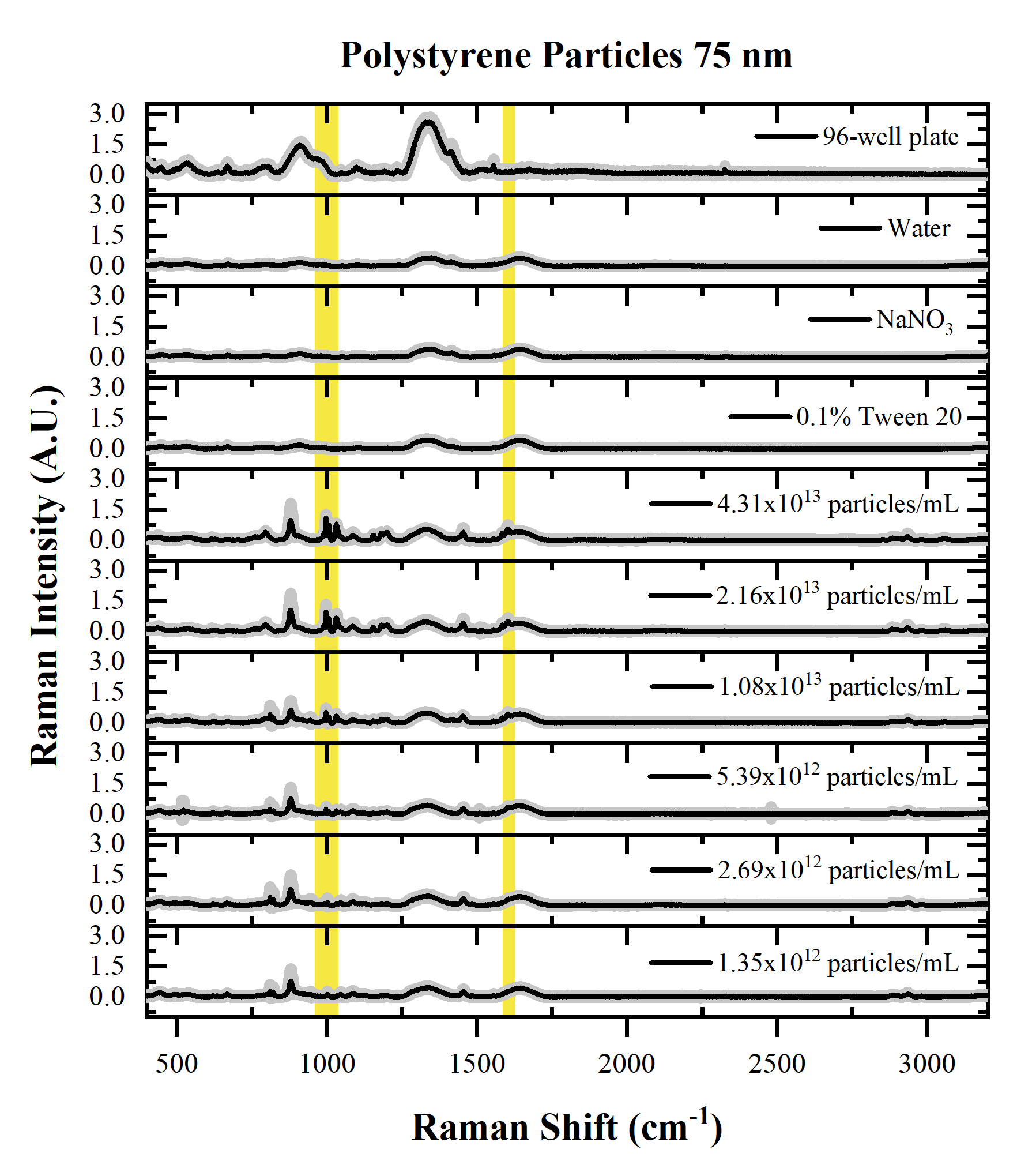

**Figure S3**: Raman spectra of 75 nm polystyrene particles. Yellow shaded area represents the area of interest of the main peaks of polystyrene at 1001 cm^-1^ (990-1017 cm^-1^), 1031 cm^-1^ (1019-1047 cm^-1^) and 1602 cm^-1^ (1592-1615 cm^-1^). Raman spectra are averaged (n=24) and gray shaded area represents the standard deviation.

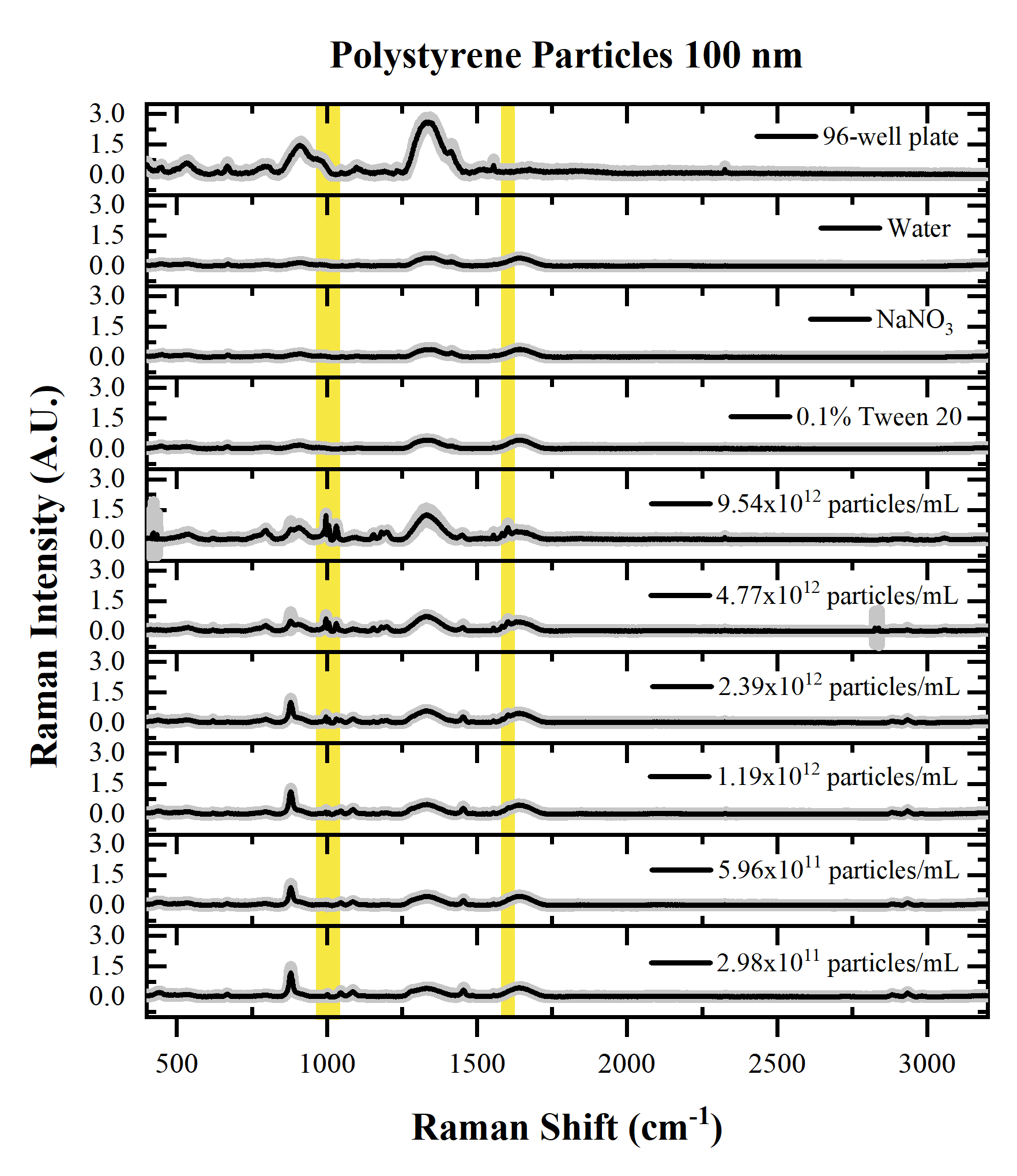

**Figure S4**: Raman spectra of 100 nm polystyrene particles. Yellow shaded area represents the area of interest of the main peaks of polystyrene at 1001 cm^-1^ (990-1017 cm^-1^), 1031 cm^-1^ (1019-1047 cm^-1^) and 1602 cm^-1^ (1592-1615 cm^-1^). Raman spectra are averaged (n=24) and gray shaded area represents the standard deviation.

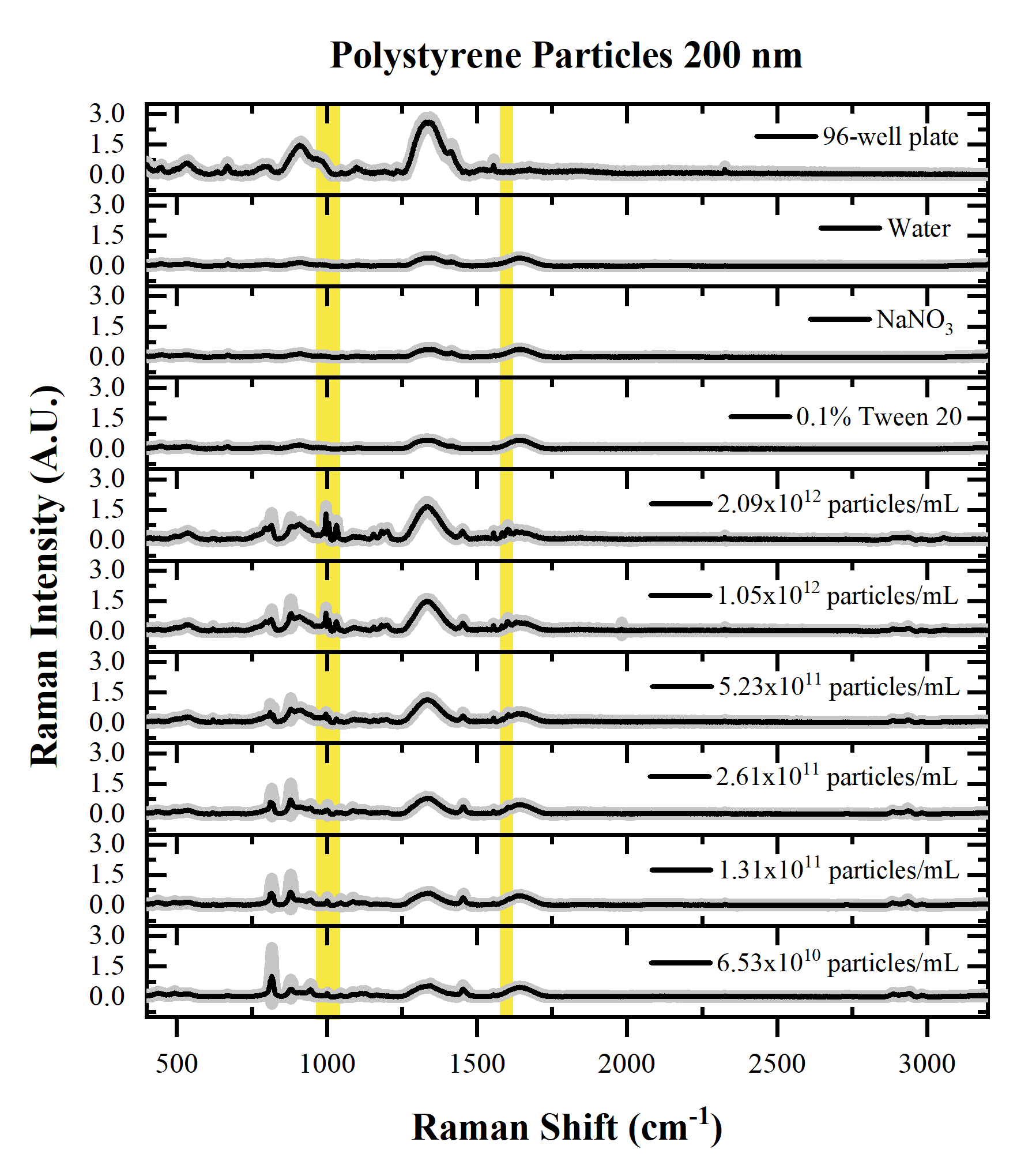

**Figure S5**: Raman spectra of 200 nm polystyrene particles. Yellow shaded area represents the area of interest of the main peaks of polystyrene at 1001 cm^-1^ (990-1017 cm^-1^), 1031 cm^-1^ (1019-1047 cm^-1^) and 1602 cm^-1^ (1592-1615 cm^-1^). Raman spectra are averaged (n=24) and gray shaded area represents the standard deviation.

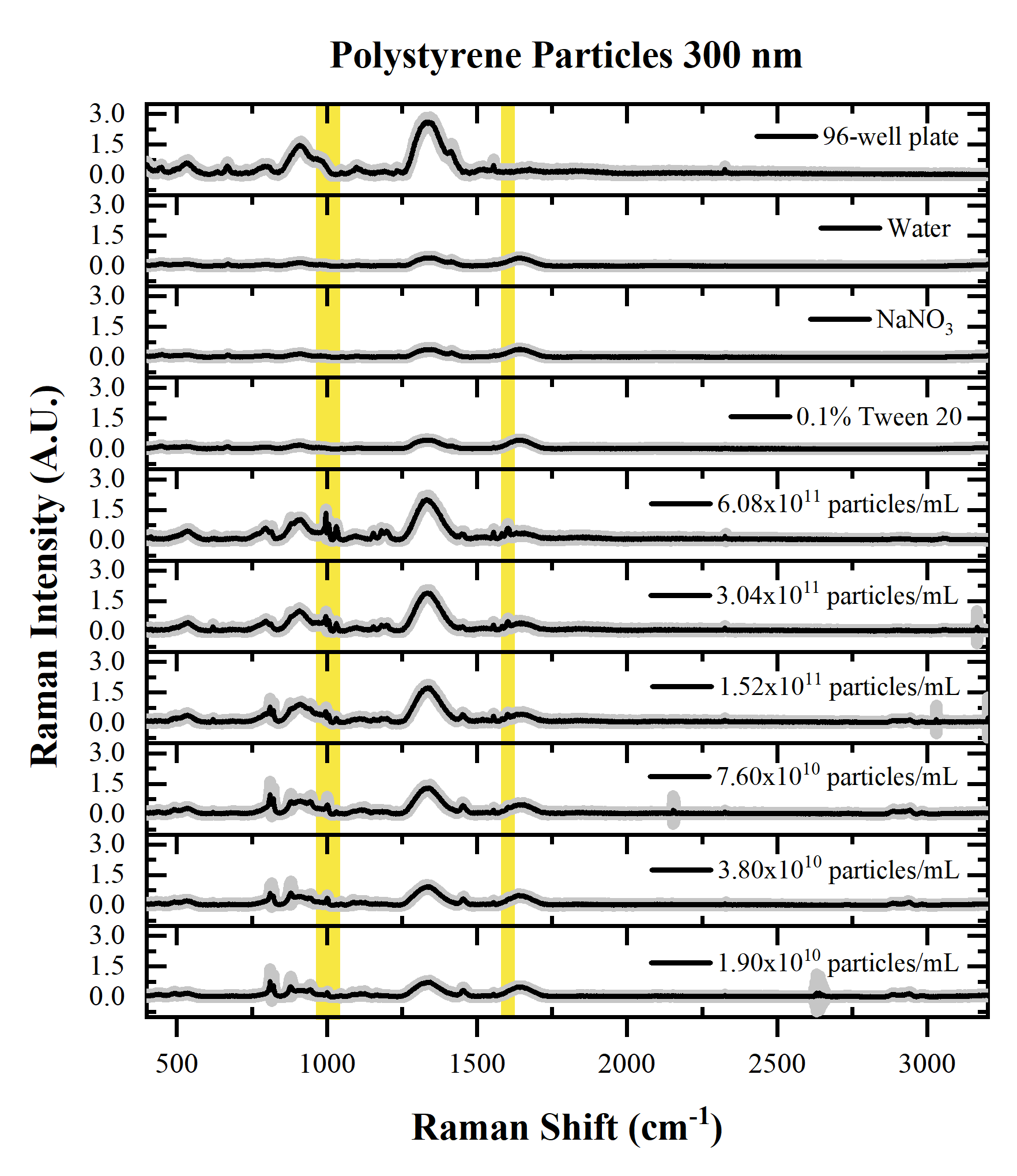

**Figure S6**: Raman spectra of 300 nm polystyrene particles. Yellow shaded area represents the area of interest of the main peaks of polystyrene at 1001 cm^-1^ (990-1017 cm^-1^), 1031 cm^-1^ (1019-1047 cm^-1^) and 1602 cm^-1^ (1592-1615 cm^-1^). Raman spectra are averaged (n=24) and gray shaded area represents the standard deviation.

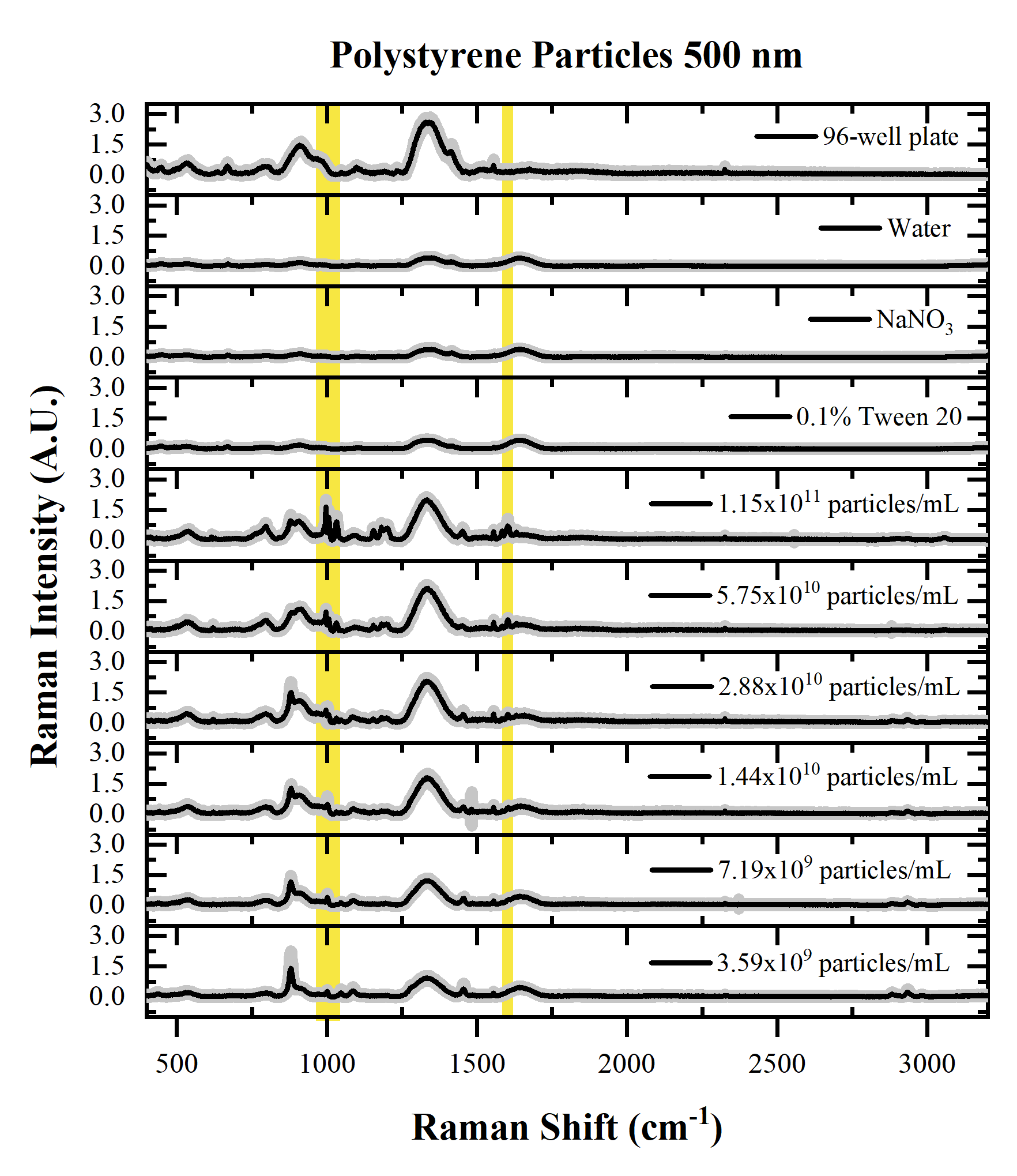

**Figure S7**: Raman spectra of 500 nm polystyrene particles. Yellow shaded area represents the area of interest of the main peaks of polystyrene at 1001 cm^-1^ (990-1017 cm^-1^), 1031 cm^-1^ (1019-1047 cm^-1^) and 1602 cm^-1^ (1592-1615 cm^-1^). Raman spectra are averaged (n=24) and gray shaded area represents the standard deviation.

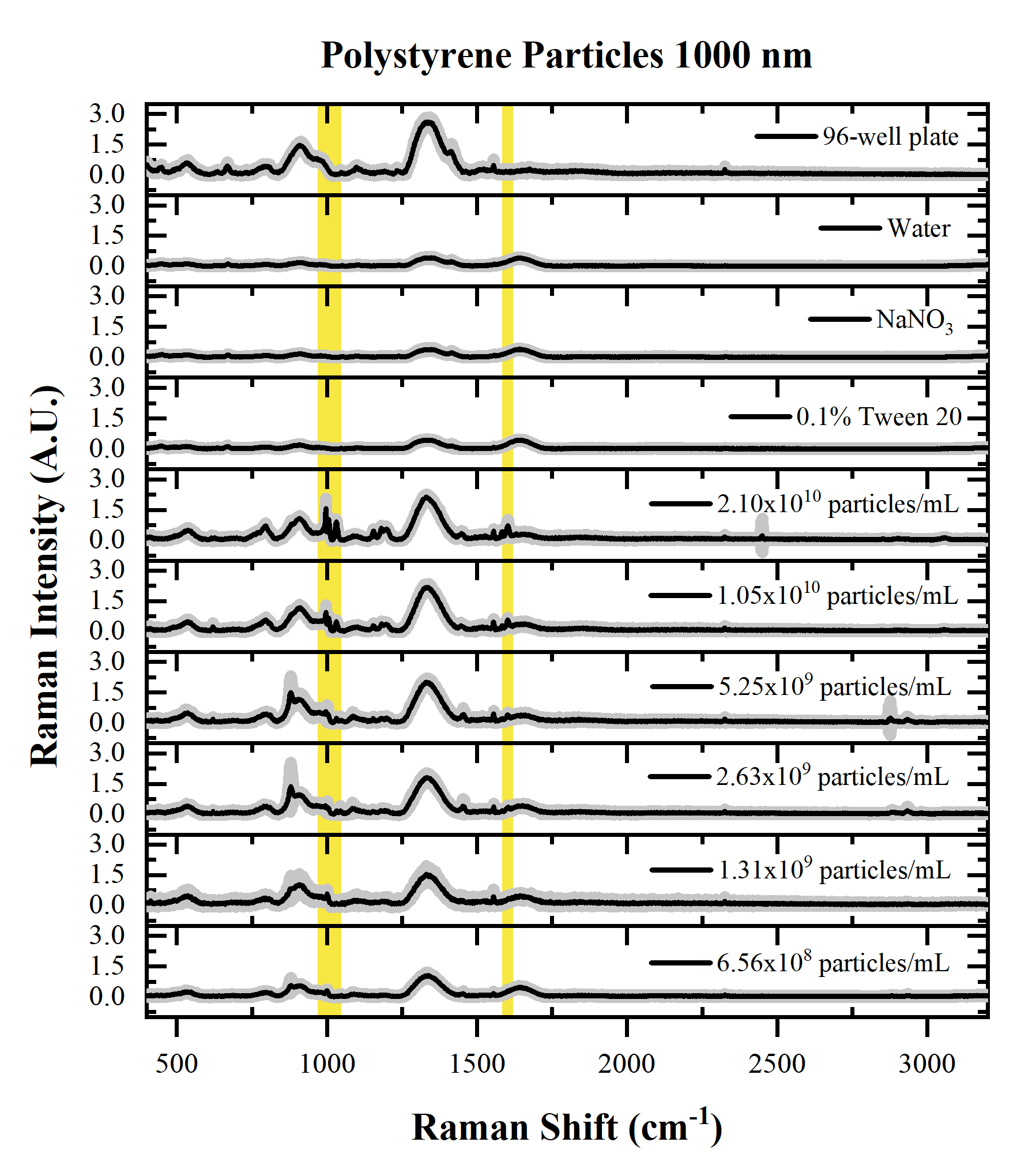

**Figure S8**: Raman spectra of 1000 nm polystyrene particles. Yellow shaded area represents the area of interest of the main peaks of polystyrene at 1001 cm^-1^ (990-1017 cm^-1^), 1031 cm^-1^ (1019-1047 cm^-1^) and 1602 cm^-1^ (1592-1615 cm^-1^). Raman spectra are averaged (n=24) and gray shaded area represents the standard deviation.

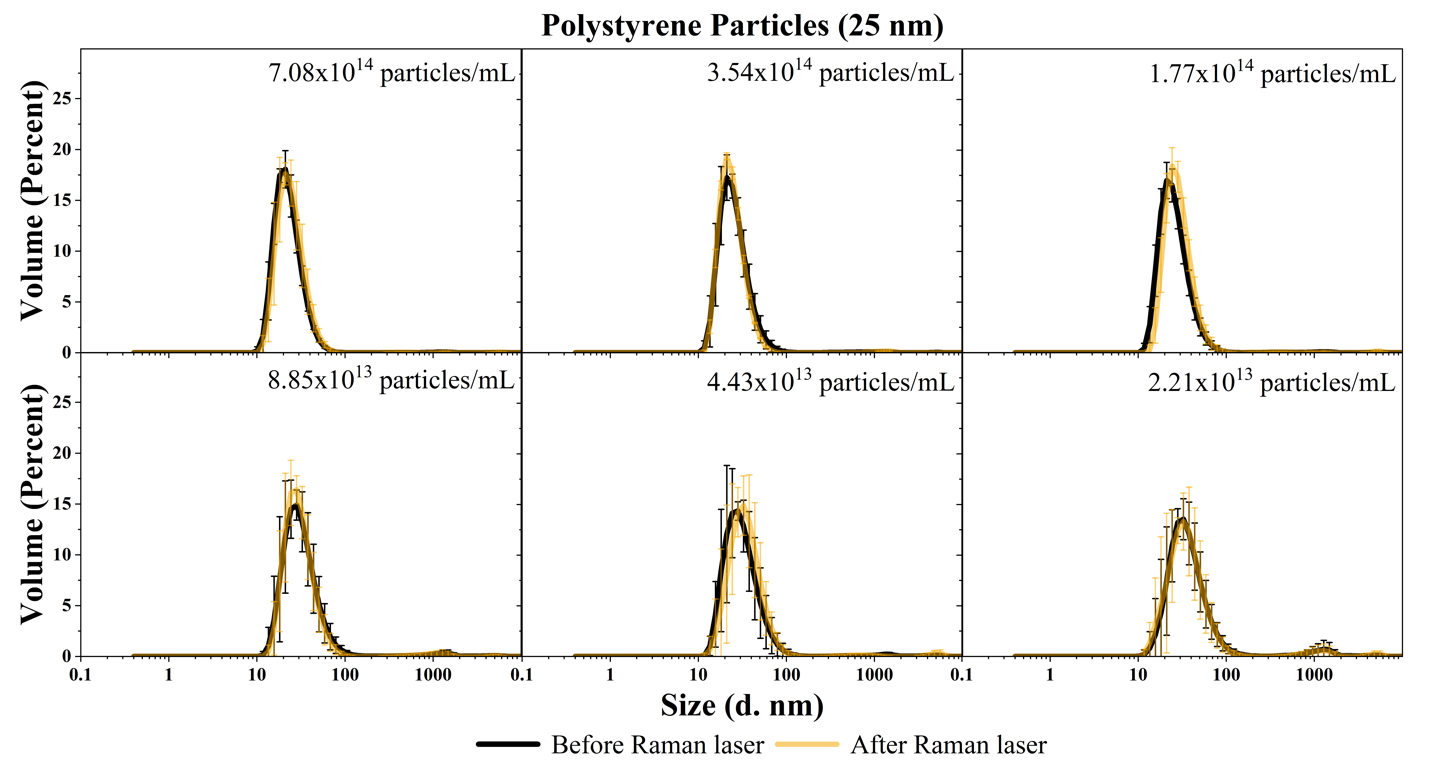

**Figure S9**: Particle size measurements of 25 nm polystyrene particles. Black line represents particle size measurements before exposure to Raman laser and orange line represents particle size measurements after exposure to Raman laser. Measurements are averaged (n=9) and error bars represent the standard deviation.

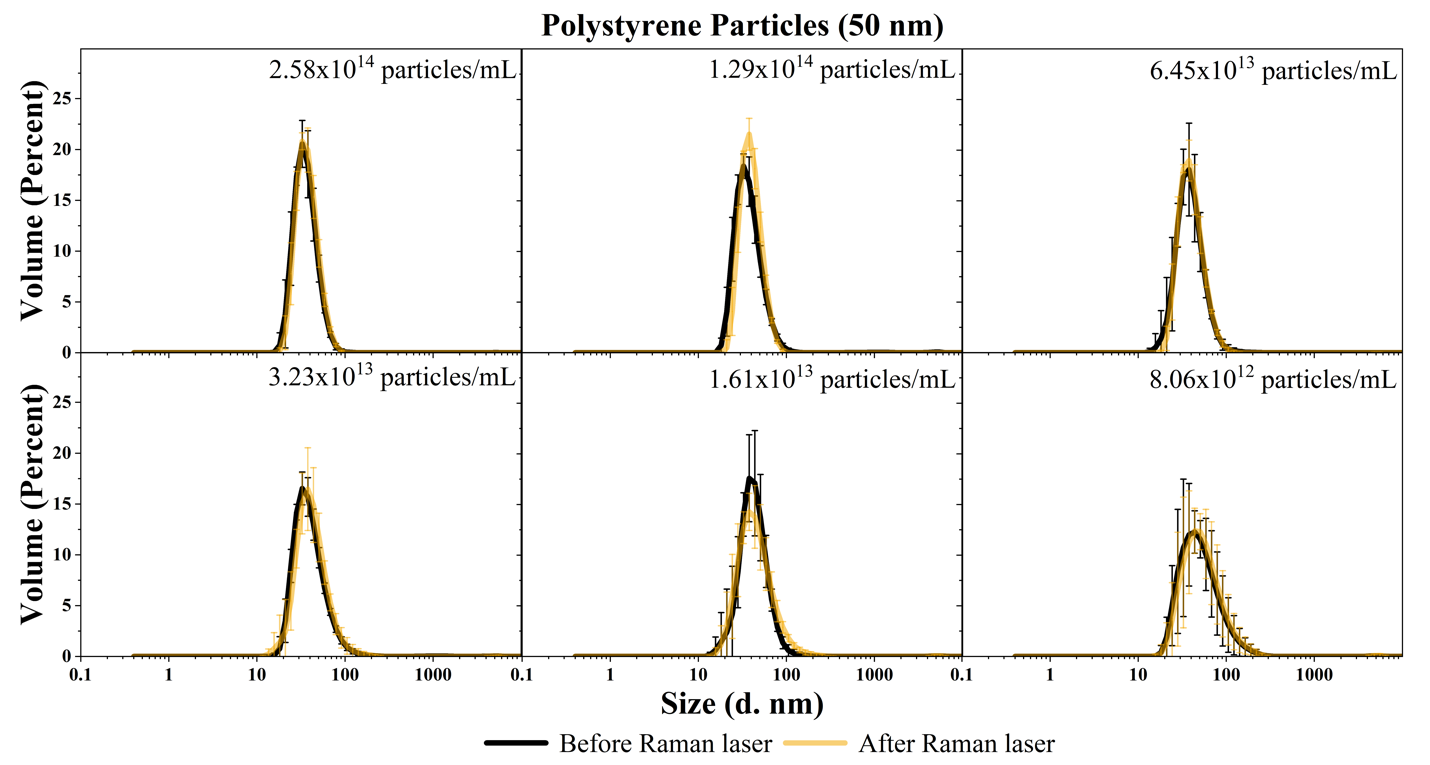

**Figure S10**: Particle size measurements of 50 nm polystyrene particles. Black line represents particle size measurements before exposure to Raman laser and orange line represents particle size measurements after exposure to Raman laser. Measurements are averaged (n=9) and error bars represent the standard deviation.

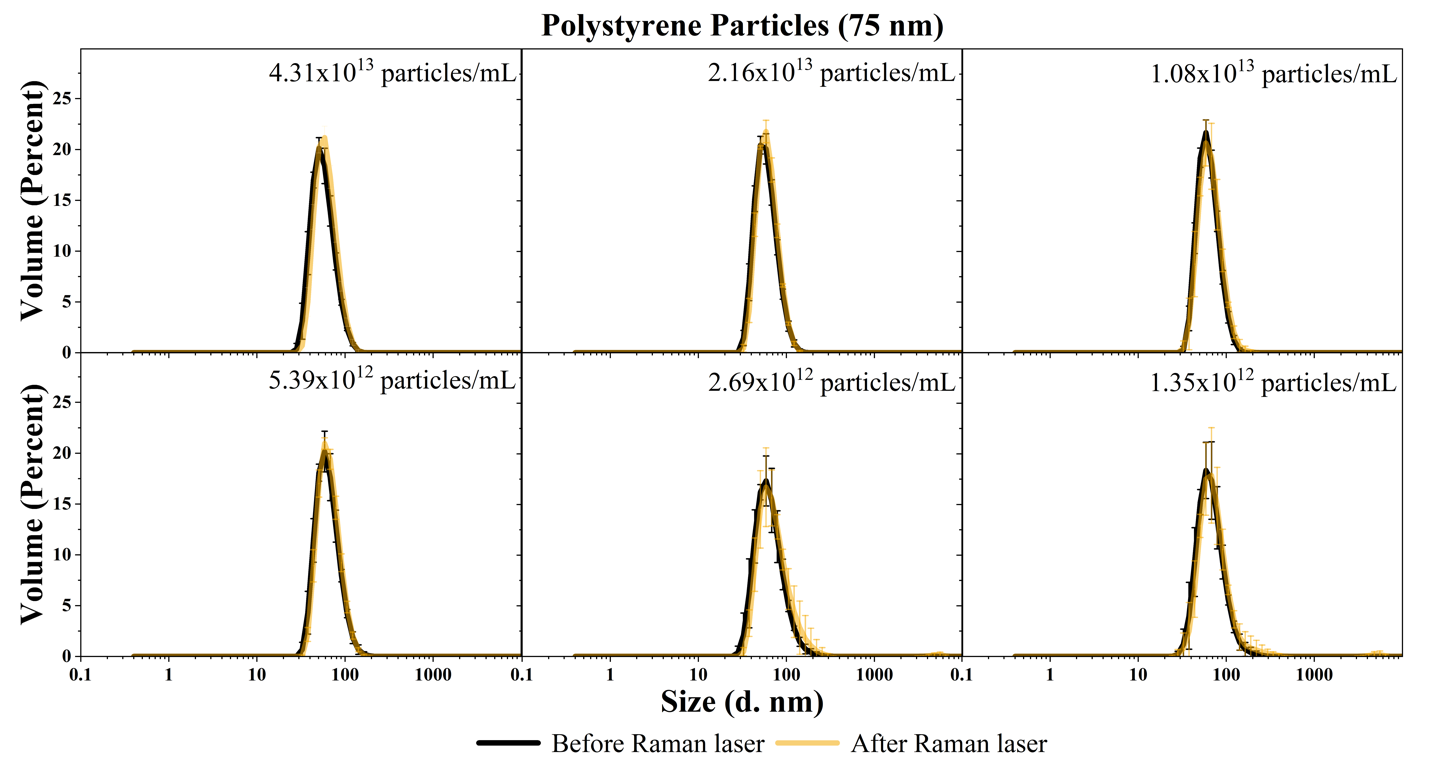

**Figure S11**: Particle size measurements of 75 nm polystyrene particles. Black line represents particle size measurements before exposure to Raman laser and orange line represents particle size measurements after exposure to Raman laser. Measurements are averaged (n=9) and error bars represent the standard deviation.

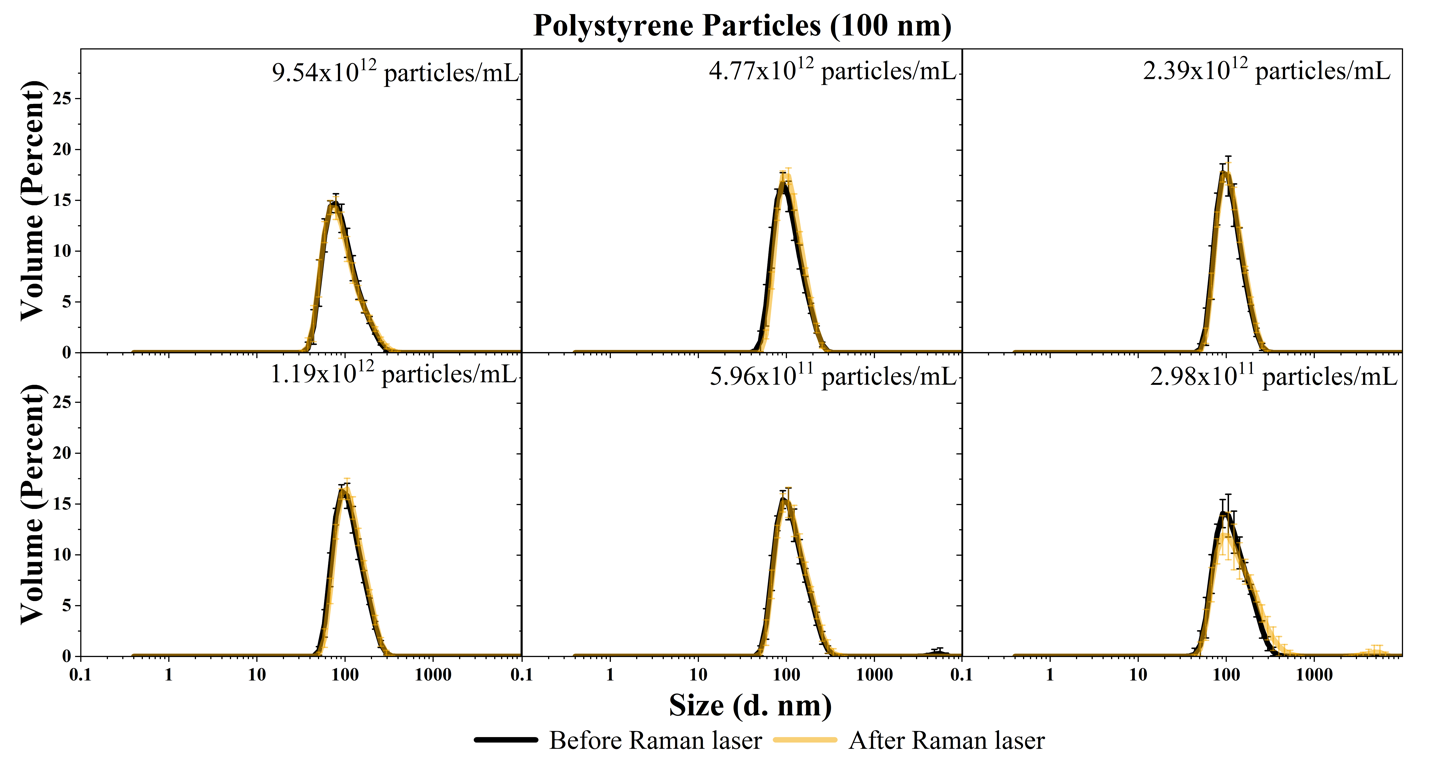

**Figure S12**: Particle size measurements of 100 nm polystyrene particles. Black line represents particle size measurements before exposure to Raman laser and orange line represents particle size measurements after exposure to Raman laser. Measurements are averaged (n=9) and error bars represent the standard deviation.

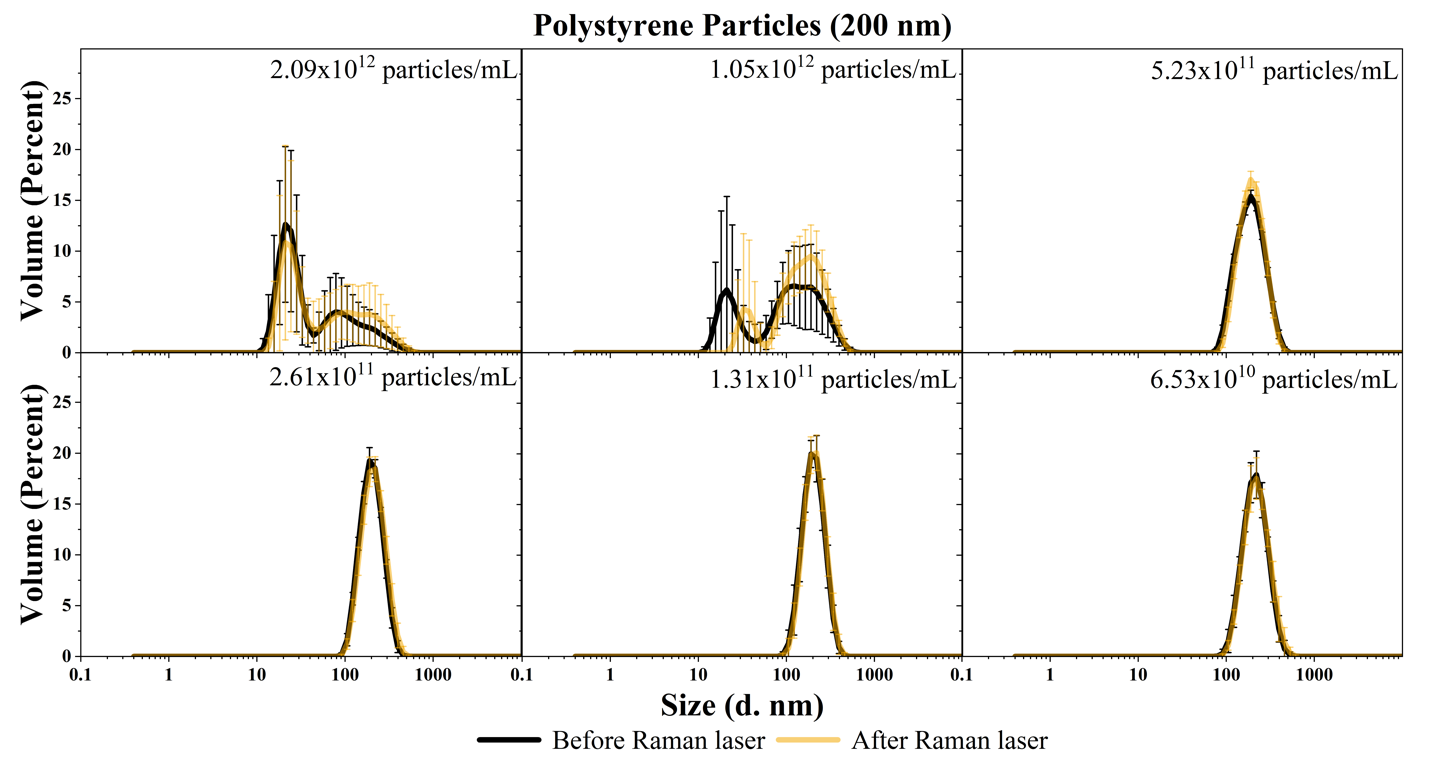

**Figure S13**: Particle size measurements of 200 nm polystyrene particles. Black line represents particle size measurements before exposure to Raman laser and orange line represents particle size measurements after exposure to Raman laser. Measurements are averaged (n=9) and error bars represent the standard deviation.

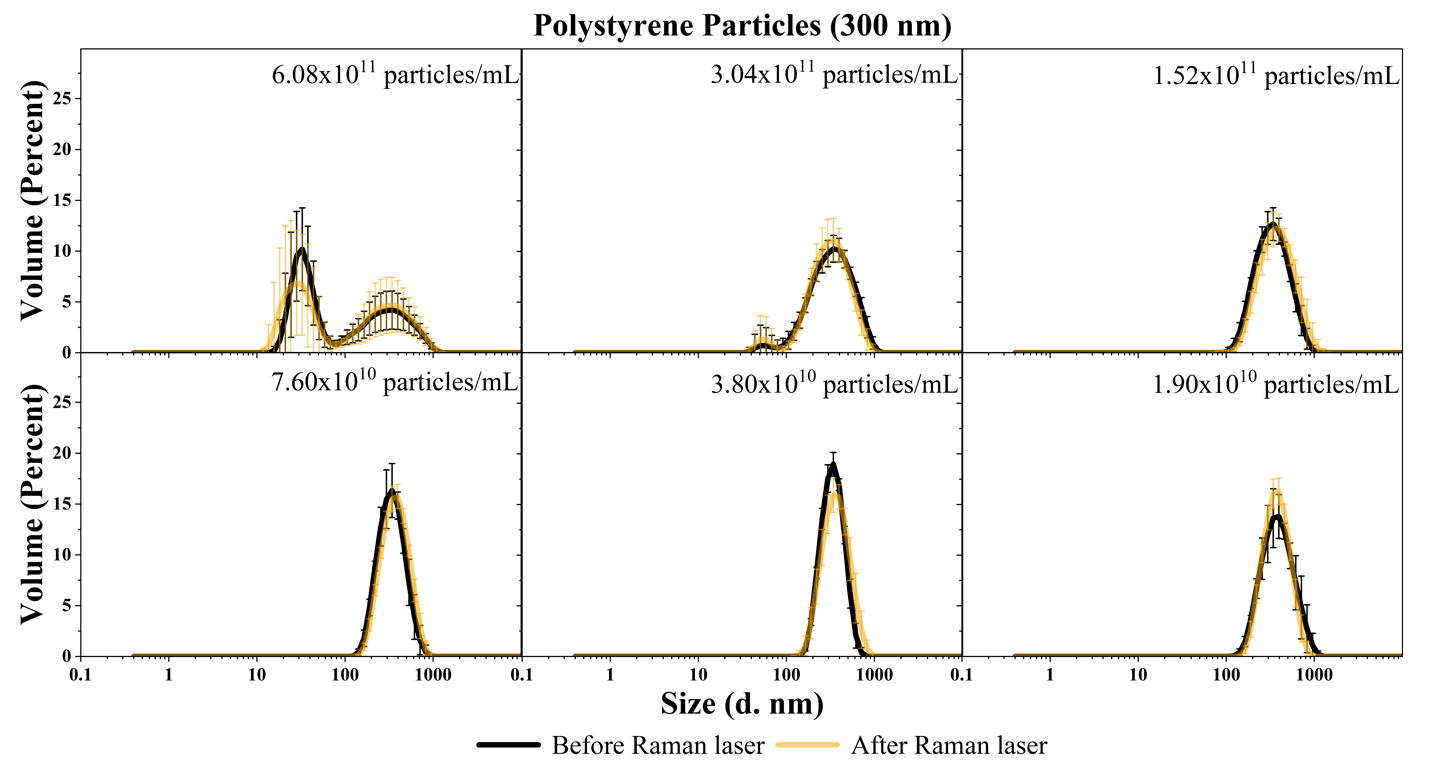

**Figure S14**: Particle size measurements of 300 nm polystyrene particles. Black line represents particle size measurements before exposure to Raman laser and orange line represents particle size measurements after exposure to Raman laser. Measurements are averaged (n=9) and error bars represent the standard deviation.

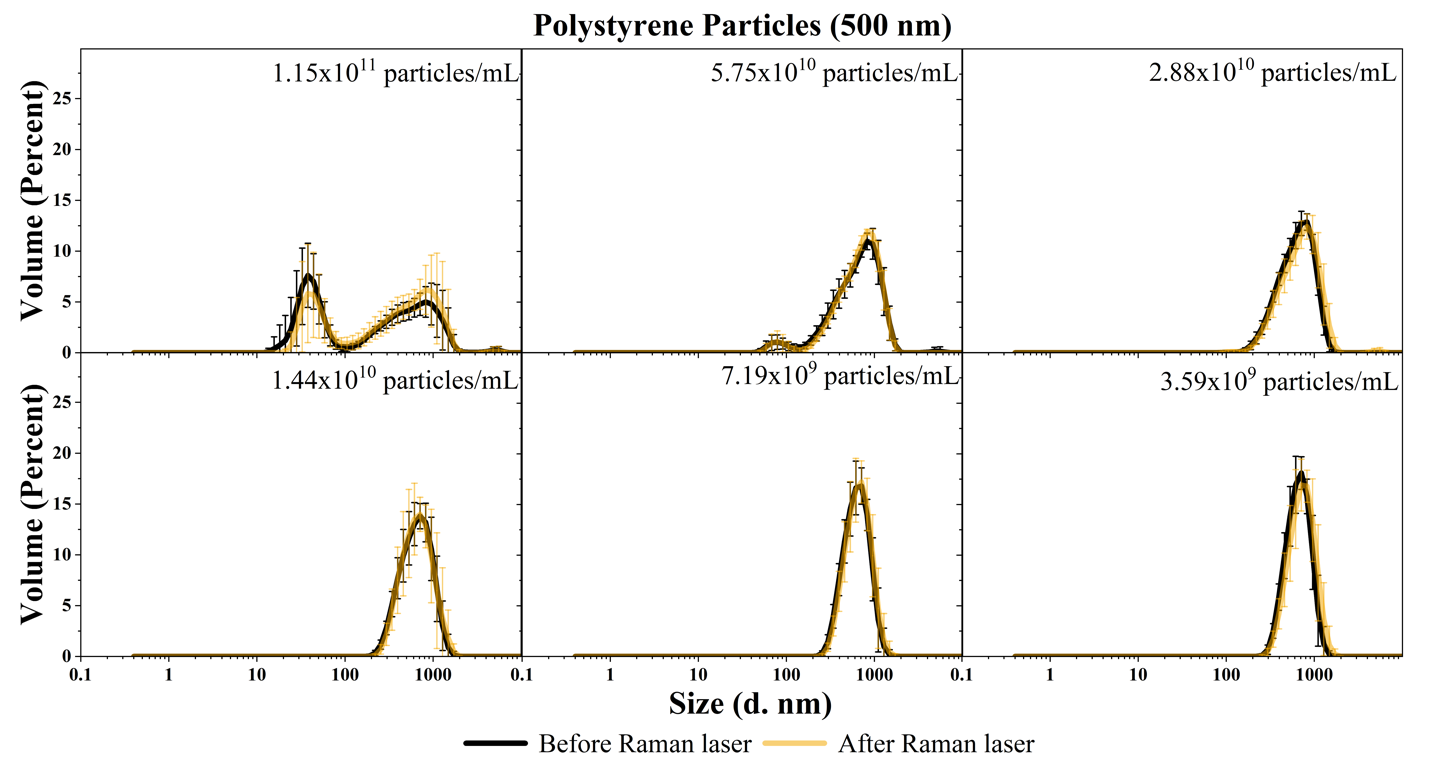

**Figure S15**: Particle size measurements of 500 nm polystyrene particles. Black line represents particle size measurements before exposure to Raman laser and orange line represents particle size measurements after exposure to Raman laser. Measurements are averaged (n=9) and error bars represent the standard deviation.

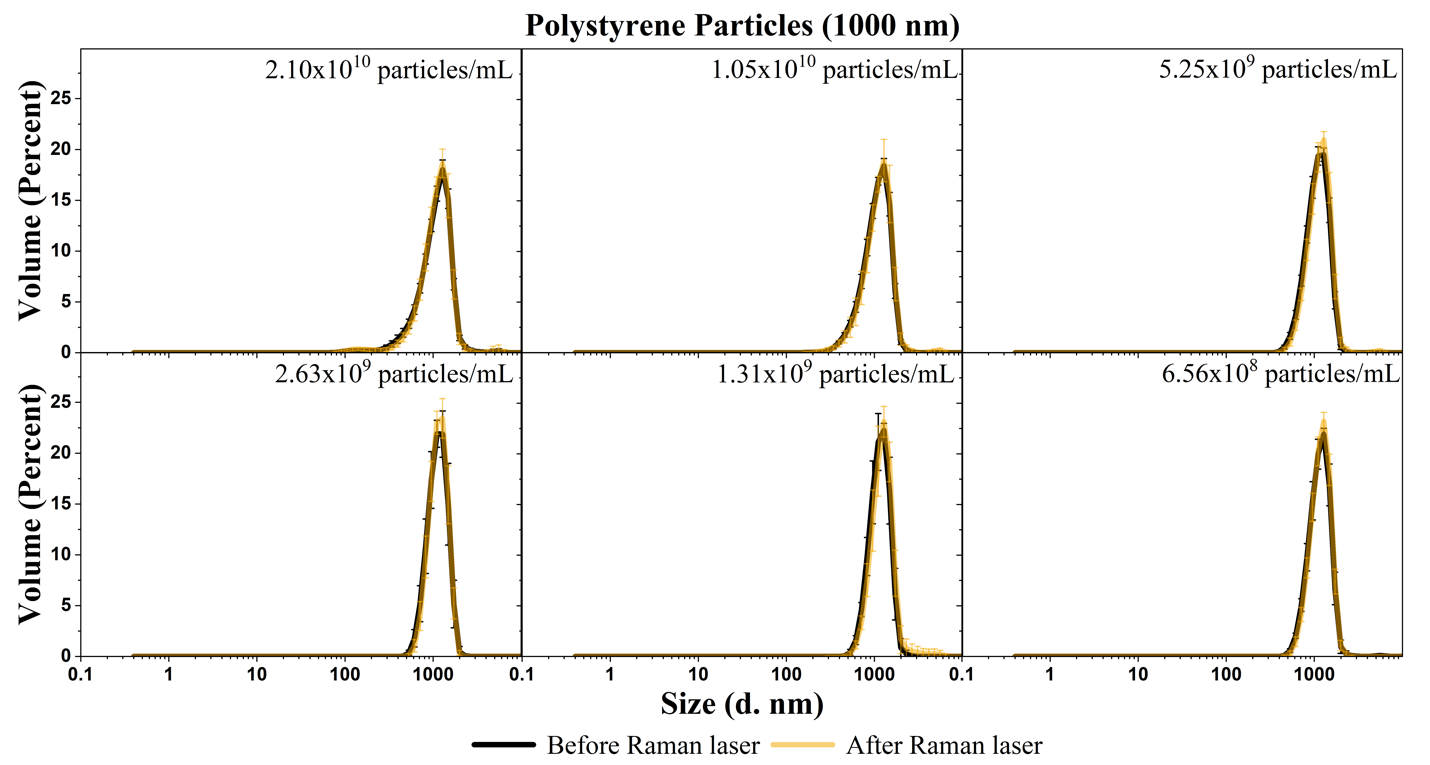

**Figure S16**: Particle size measurements of 1000 nm polystyrene particles. Black line represents particle size measurements before exposure to Raman laser and orange line represents particle size measurements after exposure to Raman laser. Measurements are averaged (n=9) and error bars represent the standard deviation.

### Supporting Tables

**Table S2**: Area under the curve (AUC) for characteristic peaks as a function of concentration, surface area, and volume.

| **Polystyrene Particles** | | | | | | |
| --- | --- | --- | --- | --- | --- | --- |
| **Size** | **Concentration** | | | **AUC (A.U. cm^-1^) for peaks at:** | | |
|  | **Concentration (particle/mL)** | **Surface area (m^2^)/mL** | **Volume (m^3^)/mL** | **1001 (cm^-1^) (990-1017 cm^-1^)** | **1031 (cm^-1^) (1019-1047 cm^-1^)** | **1602 (cm^-1^) (1592-1615 cm^-1^)** |
|  |  |  |  | n=24 ± SD | n=24 ± SD | n=24 ± SD |
| **Buffer** |  | | | 0.778 ± 0.154 | 0.237 ± 0.080 | 5.746 ± 1.261 |
| **25 nm** | 7.08x10^14^ | 1.39x10 | 5.79x10^-9^ | CCD Saturation | | |
|  | 3.54x10^14^ | 6.95x10^-1^ | 2.90x10^-9^ |  |  |  |
|  | 1.77x10^14^ | 3.48x10^-1^ | 1.45x10^-9^ |  |  |  |
|  | 8.85x10^13^ | 1.74x10^-1^ | 7.24x10^-10^ |  |  |  |
|  | 4.43x10^13^ | 8.70x10^-2^ | 3.62x10^-10^ | 1.230 ± 0.222 | 0.930 ± 0.213 | 2.711 ± 0.923 |
|  | 2.21x10^13^ | 4.34x10^-2^ | 1.81x10^-10^ | 0.791 ± 0.496 | 0.616 ±0.364 | 4.340 ± 0.955 |
| **50 nm** | 2.58x10^14^ | 2.03x10 | 1.69x10^-8^ | 7.259 ± 0.815 | 7.253 ± 1.359 | 8.872 ± 1.099 |
|  | 1.29x10^14^ | 1.01x10 | 8.44x10^-9^ | 4.342 ± 1.816 | 3.435 ± 0.559 | 7.650 ± 0.672 |
|  | 6.45x10^13^ | 5.07x10^-1^ | 4.22x10^-9^ | 2.393 ± 0.997 | 2.287 ± 1.041 | 7.114 ± 0.346 |
|  | 3.23 x10^13^ | 2.54x10^-1^ | 2.11x10^-9^ | 2.655 ± 1.250 | 1.425 ± 1.174 | 6.474 ± 0.449 |
|  | 1.61x10^13^ | 1.26x10^-1^ | 1.05x10^-9^ | 1.615 ± 0.663 | 0.960 ± 0.943 | 6.164 ± 0.413 |
|  | 8.06x10^12^ | 6.33 x10^-2^ | 5.28x10^-9^ | 1.195 ± 0.155 | 0.603 ± 0.506 | 6.106 ± 0.422 |
| **75 nm** | 4.31x10^13^ | 7.62x10^-1^ | 9.52x10^-9^ | 8.649 ± 0.475 | 9.684 ± 1.133 | 10.364 ± 1.258 |
|  | 2.16x10^13^ | 3.82x10^-1^ | 4.77x10^-9^ | 7.258 ± 2.365 | 8.754 ± 2.600 | 9.386 ± 1.288 |
|  | 1.08x10^13^ | 1.91x10^-1^ | 2.39x10^-9^ | 4.193 ± 1.327 | 4.308 ± 0.914 | 8.275 ± 0.855 |
|  | 5.39x10^12^ | 9.52x10^-1^ | 1.19x10^-9^ | 1.981 ± 0.713 | 2.549 ± 0.551 | 7.045 ± 0.617 |
|  | 2.69x10^12^ | 4.75x10^-2^ | 5.94x10^-10^ | 1.838 ± 1.313 | 1.702 ± 0.602 | 6.582 ± 0.588 |
|  | 1.35x10^12^ | 2.39x10^-2^ | 2.98x10^-10^ | 1.346 ± 0.819 | 1.216 ± 0.664 | 6.163 ± 0.451 |
| **100 nm** | 9.54x10^12^ | 3.00x10^-1^ | 5.00x10^-9^ | 9.773 ± 0.586 | 7.565 ± 0.873 | 10.770 ± 1.524 |
|  | 4.77x10^12^ | 1.50x10^-1^ | 2.50x10^-9^ | 3.779 ± 2.192 | 3.539 ± 1.654 | 9.047 ± 0.697 |
|  | 2.39x10^12^ | 7.51x10^-1^ | 1.25x10^-9^ | 2.509 ± 0.232 | 2.929 ± 0.187 | 7.784 ± 0.597 |
|  | 1.19x10^12^ | 3.74x10^-2^ | 6.23x10^-10^ | 1.182 ± 0.149 | 2.067 ± 0.225 | 6.773 ± 0.299 |
|  | 5.96 x10^11^ | 1.87x10^-2^ | 3.12x10^-10^ | 0.775 ± 0.384 | 1.377 ± 0.111 | 6.418 ± 0.352 |
|  | 2.98x10^11^ | 9.36x10^-3^ | 1.56x10^-10^ | 1.107 ± 0.145 | 1.750 ± 0.576 | 6.255 ± 0.519 |

Continuation of Table S1

| **Polystyrene Particles** | | | | | | |
| --- | --- | --- | --- | --- | --- | --- |
| **Size** | **Concentration** | | | **AUC for peaks at:** | | |
|  | **Concentration (particle/mL)** | **Surface area (m^2^)/mL** | **Volume (m^3^)/mL** | **1001 (cm^-1^) (990-1017 cm^-1^)** | **1031 (cm^-1^) (1019-1047 cm^-1^)** | **1602 (cm^-1^) (1592-1615 cm^-1^)** |
|  |  |  |  | n=24 ± SD | n=24 ± SD | n=24 ± SD |
| **200 nm** | 2.09x10^12^ | 2.63x10^-1^ | 8.75x10^-9^ | 10.951 ± 2.286 | 8.024 ± 2.552 | 10.008 ± 1.341 |
|  | 1.05x10^12^ | 1.31x10^-1^ | 4.38x10^-9^ | 8.143 ± 1.669 | 5.456 ± 0.902 | 9.005 ± 1.262 |
|  | 5.23x10^11^ | 6.57x10^-2^ | 2.19x10^-9^ | 5.328 ± 1.428 | 2.402 ± 0.433 | 7.604 ± 0.719 |
|  | 2.61x10^11^ | 3.28x10^-2^ | 1.09x10^-9^ | 3.196 ± 1.832 | 1.796 ± 0.960 | 7.178 ± 0.525 |
|  | 1.31x10^11^ | 1.64x10^-2^ | 5.47x10^-10^ | 2.426 ± 1.191 | 1.572 ± 1.432 | 6.684 ± 0.363 |
|  | 6.53x10^10^ | 8.21x10^-3^ | 2.74x10^-10^ | 1.754 ± 0.335 | 0.796 ± 0.659 | 6.500 ± 0.445 |
| **300 nm** | 6.08x10^11^ | 1.72x10^-1^ | 8.60x10^-9^ | 12.024 ± 1.468 | 7.328 ± 0.803 | 9.838 ± 1.434 |
|  | 3.04x10^11^ | 8.60x10^-2^ | 4.30x10^-9^ | 9.330 ± 2.354 | 4.183 ± 0.484 | 8.113 ± 0.881 |
|  | 1.52x10^11^ | 4.30x10^-2^ | 2.15x10^-9^ | 7.364 ± 1.133 | 2.590 ± 0.371 | 7.460 ± 0.725 |
|  | 7.60x10^10^ | 2.15x10^-2^ | 1.07x10^-9^ | 6.314 ± 2.474 | 1.466 ± 0.272 | 6.987 ± 0.577 |
|  | 3.80x10^10^ | 1.07x10^-2^ | 5.37x10^-10^ | 3.961 ± 0.835 | 1.015 ± 0.497 | 6.604 ± 0.430 |
|  | 1.90x10^10^ | 5.37x10^-3^ | 2.69x10^-10^ | 2.373 ± 0.490 | 0.713 ± 0.464 | 6.578 ± 0.519 |
| **500 nm** | 1.15x10^11^ | 9.03^X^10^-2^ | 7.53x10^-9^ | 13.746 ± 2.445 | 10.401 ± 3.067 | 11.348 ± 2.437 |
|  | 5.75x10^10^ | 4.52x10^-2^ | 3.76x10^-9^ | 10.199 ± 1.098 | 4.650 ± 0.652 | 8.581 ± 0.701 |
|  | 2.88x10^10^ | 2.26x10^-2^ | 1.88x10^-9^ | 8.571 ± 2.862 | 3.232 ± 0.908 | 6.916 ± 0.702 |
|  | 1.44x10^10^ | 1.13x10^-2^ | 9.41x10^-10^ | 7.413 ± 2.863 | 1.837 ± 0.385 | 6.223 ± 0.844 |
|  | 7.19x10^9^ | 5.65x10^-3^ | 4.70x10^-10^ | 4.574 ± 1.211 | 1.256 ± 0.247 | 5.927 ± 0.652 |
|  | 3.59x10^9^ | 2.82x10^-3^ | 2.35x10^-10^ | 2.951 ± 0.517 | 1.735 ± 1.085 | 6.025 ± 0.505 |
| **1000 nm** | 2.10x10^10^ | 6.60x10^-2^ | 1.10x10^-8^ | 14.232 ± 1.599 | 9.517 ± 1.037 | 10.743 ± 1.882 |
|  | 1.05x10^10^ | 3.30x10^-2^ | 5.50x10^-9^ | 10.869 ± 1.161 | 5.225 ± 0.572 | 8.759 ± 0.908 |
|  | 5.25x10^9^ | 1.65x10^-2^ | 2.75x10^-9^ | 8.688 ± 3.332 | 3.107 ± 0.887 | 6.890 ± 0.993 |
|  | 2.63x10^9^ | 8.25x10^-3^ | 1.37x10^-9^ | 6.577 ± 2.167 | 2.631 ± 1.447 | 6.489 ± 0.615 |
|  | 1.31x10^9^ | 4.12x10^-3^ | 6.87x10^-10^ | 8.242 ± 2.825 | 2.019 ± 1.470 | 7.045 ± 1.646 |
|  | 6.56x10^8^ | 2.06x10^-3^ | 3.44x10^-10^ | 4.031 ± 0.533 | 0.766 ± 0.333 | 6.376 ± 0.679 |

**Table S3:** Estimated LOD ranges for polystyrene particles 50 nm-1000 nm.

| Particle Size (nm) | LOD_min_ (particle/mL) | LOD_max_ (particle/mL) |
| --- | --- | --- |
| 50 | 7.47 x 10^12^ | 7.64 x 10^12^ |
| 75 | 1.22 x 10^12^ | 1.24 x 10^12^ |
| 100 | 2.74 x 10^11^ | 2.91 x 10^11^ |
| 200 | 5.99 x 10^10^ | 6.04 x 10^10^ |
| 300 | 1.74 x 10^10^ | 1.78 x 10^10^ |
| 500 | 3.34 x 10^9^ | 3.37 x 10^9^ |
| 1000 | 6.09 x 10^8^ | 6.24 x 10^8^ |

### Partial Least Squares (PLS) Regression

RMSEP: 0.058049475251639254

VIP Scores:

- Size: 1.3915549915893195
- Concentration: 1.329223969382576
- Day: 0.5542135937849549
- Replicates: 0.40575397197537405

### Limit of Detection

Sensitivity: 1.1717677827987376

Size: 50 nm

Variance of X: 7.030037994605294E+24

Variance of y_cal: 0.0008393783040921155

h_0min_: 3.0888445373716866E-06

h_0max_: 0.04654617518528472

LOD_min_: 7.467099E+12

LOD_max_: 7.638893E+12

Size: 75 nm

Variance of X: 1.8730357264365508E+23

Variance of y_cal: 0.00036574617644881906

h_0min_: 3.481019248121668E-06

h_0max_: 0.027020974914847458

LOD_min_: 1.218839E+12

LOD_max_: 1.235194E+12

Size: 100 nm

Variance of X: 9.457526284532199E+21

Variance of y_cal: 0.0010678995338423702

h_0min_: 4.73071226403998E-06

h_0max_: 0.12908953084048586

LOD_min_: 2.738812E+11

LOD_max_: 2.910217E+11

Size: 200 nm

Variance of X: 4.527833585440157E+20

Variance of y_cal: 0.0008199387889638149

h_0min_: 4.753914576206163E-06

h_0max_: 0.017056431161975538

LOD_min_: 5.992646E+10

LOD_max_: 6.043522E+10

Size: 300 nm

Variance of X: 3.812109402550231E+19

Variance of y_cal: 0.0013358852815019528

h_0min_: 5.96686156538727E-06

h_0max_: 0.04489558860414215

LOD_min_: 1.738827E+10

LOD_max_: 1.777426E+10

Size: 500 nm

Variance of X: 1.4086780918943972E+18

Variance of y_cal: 0.0011530429034752778

h_0min_: 6.1010863470212805E-06

h_0max_: 0.018636157097288227

LOD_min_: 3.342563E+09

LOD_max_: 3.373555E+09

Size: 1000 nm

Variance of X: 4.675557774057551E+16

Variance of y_cal: 0.0028244984025060783

h_0min_: 6.494905145385191E-06

h_0max_: 0.048340791538837774

LOD_min_: 6.089624E+08

LOD_max_: 6.235056E+08

### References

(1)	Wheaton, S.; Gelfand, R. M.; Gordon, R. Probing the Raman-Active Acoustic Vibrations of Nanoparticles with Extraordinary Spectral Resolution. *Nat. Photonics* **2015**, *9* (1), 68–72. https://doi.org/10.1038/nphoton.2014.283.

(2)	Pollard, M. R.; Sparnacci, K.; Wacker, L. J.; Kerdoncuff, H. Polymer Nanoparticle Identification and Concentration Measurement Using Fiber-Enhanced Raman Spectroscopy. *Chemosensors* **2020**, *8* (1), 21. https://doi.org/10.3390/chemosensors8010021.

(3)	Meyer-Kirschner, J.; Mitsos, A.; Viell, J. Polymer Particle Sizing from Raman Spectra by Regression of Hard Model Parameters. *J. Raman Spectrosc.* **2018**, *49* (8), 1402–1411. https://doi.org/10.1002/jrs.5387.

(4)	Dorney, J.; Bonnier, F.; Garcia, A.; Casey, A.; Chambers, G.; J. Byrne, H. Identifying and Localizing Intracellular Nanoparticles Using Raman Spectroscopy. *Analyst* **2012**, *137* (5), 1111–1119. https://doi.org/10.1039/C2AN15977E.

(5)	Kerman, S.; Chang, C.; Li, Y.; Lagae, L.; Stakenborg, T.; Dorpe, P. V. Raman Spectroscopy and Optical Trapping of 20 Nm Polystyrene Particles in Plasmonic Nanopores; 2014; Vol. 9126, p 912612. https://doi.org/10.1117/12.2052609.

(6)	Kerman, S.; Chen, C.; Li, Y.; Roy, W. V.; Lagae, L.; Dorpe, P. V. Raman Fingerprinting of Single Dielectric Nanoparticles in Plasmonic Nanopores. *Nanoscale* **2015**, *7* (44), 18612–18618. https://doi.org/10.1039/C5NR05341B.

(7)	Bridges, T. E.; Houlne, M. P.; Harris, J. M. Spatially Resolved Analysis of Small Particles by Confocal Raman Microscopy: Depth Profiling and Optical Trapping. *Anal. Chem.* **2004**, *76* (3), 576–584. https://doi.org/10.1021/ac034969s.

1. This value was provided by the corresponding author of this paper by email communication.
